# Inflammatory CD4/CD8 double positive human T cells arise from reactive CD8 T cells and are sufficient to mediate GVHD pathology

**DOI:** 10.1101/2022.01.11.475845

**Authors:** Nicholas J. Hess, David P. Turicek, Jeremiah Riendeau, Sean J. McIlwain, Emmanuel Contreras Guzman, Kalyan Nadiminti, Amy Hudson, Natalie S. Callander, Melissa C. Skala, Jenny E. Gumperz, Peiman Hematti, Christian M. Capitini

## Abstract

An important paradigm in allogeneic hematopoietic cell transplantations (allo-HCTs) is the prevention of graft-vs-host disease (GVHD) while preserving the graft-vs-leukemia (GVL) activity of donor T cells. From an observational clinical study of adult allo-HCT recipients, we identified a CD4+/CD8+ double positive T cell (DPT) population, not present in starting grafts, whose presence was predictive of ≥ grade 2 GVHD. Using an established xenogeneic transplant model, we reveal that the DPT population develop from antigen stimulated CD8 T cells which become transcriptionally, metabolically and phenotypically distinct from single-positive CD4 and CD8 T cells. Isolated DPTs were sufficient to mediate xeno-GVHD pathology when re-transplanted into naive mice but provided no survival benefit when mice were challenged with a human B-ALL cell line. Overall, this study reveals human DPTs as a T cell population directly involved with GVHD pathology.

**One Sentence Summary:** Human CD4+/CD8+ double positive T cells (DPTs) mediate xenogeneic GVHD but possess limited GVL activity.

## Introduction

Graft-vs-host disease (GVHD) and relapse remain the primary complications following allogeneic hematopoietic cell transplantation (allo-HCT)^1–3^. While the field has made substantial strides in reducing GVHD, current GVHD prophylaxis drug regimens target the entire T cell population, potentially hindering the efficacy of graft-vs-leukemia (GVL) activity mediated by donor T cells^4–8^. Thus, identifying predictive biomarkers and delineating specific cellular mechanism(s) leading to GVHD pathology versus GVL activity has remained a top priority for the field.

Unfortunately, separating the graft-vs-host (GVH) response of donor T cells from their GVL activity has proven difficult as both responses are derived from allogeneic antigen stimulation^7–9^. Nevertheless, several studies have investigated the role of specific HLA mismatches on the development of GVHD and GVL activity^10–13^. These studies have highlighted the importance of permissive and non-permissive HLA mismatches on GVHD development and non-relapse mortality, but correlating them with GVL activity has been met with limited success^10–14^. Furthermore, the use of haploidentical transplantations with post-transplant cyclophosphamide has shown that HLA matching of the donor and recipient may not play a dominant role in separating GVH and GVL activity^15,16^.

While pre-transplant prognostic variables such as HLA-matching, conditioning regimens and graft source all influence allo-HCT outcomes, these population-based variables are not able to predict individual patient outcomes^17–19^. Several post-transplant predictive variables have been discovered including sST2, REG3α, elafin and others^20–26^. These highly promising plasma-based biomarkers are generally released due to host damage and have shown efficacy in predicting the development of steroid-resistant GVHD and non-relapse mortality (NRM) either individually or combined into a predictive algorithm^20–26^. To date though, these biomarkers have not been able to predict an allo-HCT recipients’ chance of relapse as they measure the degree of host damage rather than donor T cell alloreactivity.

While all the validated GVHD biomarkers to date have been plasma-based, the direct investigation of T cell subsets as biomarkers has been another area of active investigation^27–30^. Both CD4 and CD8 T cells can respond to allogeneic antigens, but while CD8 T cell may be directly responsible for cellular cytotoxicity, CD4 T cells have been the primary focus of the field^31–33^. The CD4 population is unique in its capacity to polarize into a variety of different subsets, based on the cytokine environment it finds itself, to drive an inflammatory response^34–40^. In the case of GVHD, TH_1_ and TH_17/22_ cells have been shown to be directly involved in driving the pathogenesis of gastrointestinal, liver and skin pathology^35,41,42^. In contrast, the CD8 T cell population is thought to be more conserved in their role as cytotoxic effectors licensed by CD4 T cells. The discovery of CD8 regulatory T cells and Tc17 cells in murine GVHD though suggests CD8 T cells may also be able to polarize into different subsets similar to the CD4 population^43–45^. T cell specific GVHD prophylaxis regimens have been (e.g., ATG) and are currently (e.g., abatacept) promising therapeutics but the next generation of T cell-targeting GVHD prophylaxis regimens will need to precisely target GVHD-specific T cell populations.

Surprisingly, there have been numerous studies in the past two decades that have observed the presence of a T cell population co-expressing both the CD4 and CD8 lineage markers. Their presence has been identified in a range of human chronic inflammatory diseases including most recently in GVHD^46–56^. While historically thought to be the result of thymocytes escaping the thymus, recent observational studies suggest CD4+/CD8+ double positive T cells (DPTs) are in fact a mature human T cell population that develops in the periphery in response to antigenic stimulation^57,58^.

In this study, we identified DPTs as a predictive biomarker of GVHD in a 35-patient observational study of allo-HCT recipients at a single institution. Using a xenogeneic transplant model system to directly investigate human T cell biology, we further investigated the functional relevance of DPTs in GVHD pathogenesis^31,59^. We identify the origin of DPTs, which are not present in healthy human grafts, as the CD8 T cell population which arises due to TCR-dependent activation signals. We also provide evidence that the transcriptional, metabolic, and phenotypically distinct DPTs are sufficient to mediate xeno-GVHD pathology but provide no survival advantage when mice are also challenged with a human B-ALL malignancy. Overall, this study provides the first comprehensive analysis of the origin, effector functions and clinical significance of a human DPT population.

### Results

#### Increased frequencies of CD4+/CD8+ double positive T cells (DPTs) are predictive of GVHD in an observational clinical cohort of allo-HCT recipients

To determine if the direct measurement of T cells in allo-HCT recipients post-transplant can serve as predictive biomarkers of relapse and/or GVHD, we collected 417 blood samples from 35 adult allo-HCT recipients over their first 100 days post-transplant (Fig 1A). Patients were included in the observational study irrespective of their conditioning regimen, donor HLA matching, graft source or GVHD prophylaxis regimen. The cohort is heavily skewed to patients receiving G-CSF mobilized peripheral blood and a cyclophosphamide based GVHD prophylaxis regimen as both conditions are standard at our clinic (Supplemental Table 1). An average of 12 samples were collected per patient and a total of 25 samples per collection interval (~every 4-6 days) (Supplemental Fig 1). Additional information on the stratification of this cohort can be found in the Materials and Methods. Samples were processed within seven days of collection and never frozen. Antigen-experienced T cells were gated based on CD3 and CD45RO expression to exclude non-reactive donor T cells and naïve T cells that are present due to de novo hematopoiesis (Supplemental Fig 2).

**Figure 1.**
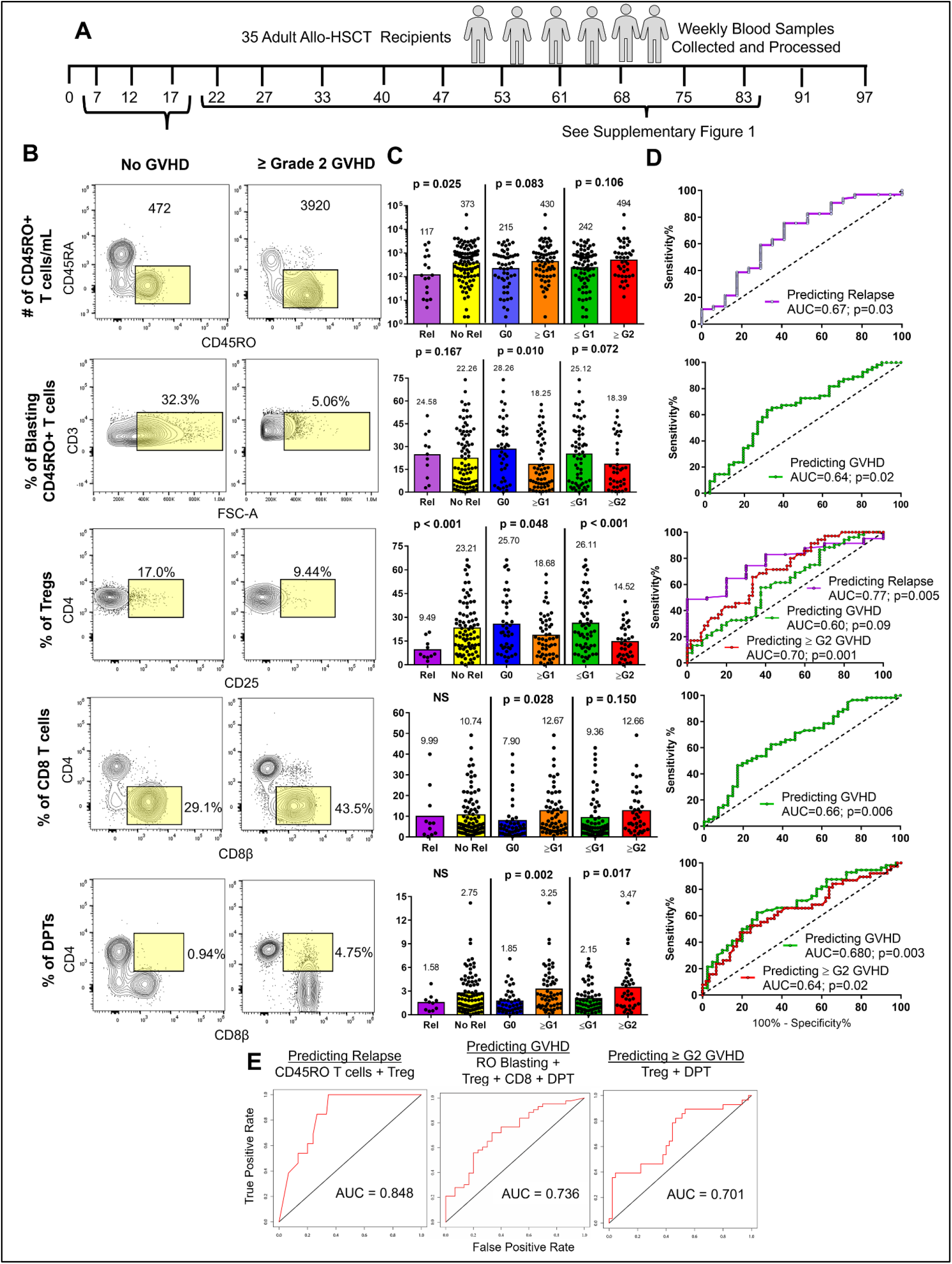
Post-transplant T cell metrics are predictive of allo-HCT recipient disease outcomes. (A) Schematic depicting the study outline. 35 allo-HCT recipients had weekly blood samples collected and processed without freezing. (B) Dot plots taken from five concatenated allo-HCT recipient samples who did not develop GVHD and those that developed ≥ grade 2 GVHD. Yellow box indicates gating strategy for each T cell metric. Samples collected between the day 7 and 17 collection intervals were used for the analysis in (C-E) while the analysis for samples collected between day 22 and 83 can be found in Supplemental Fig 3. (C) Each T cell metric was analyzed in three binary tests comparing two allo-HCT outcomes. Mean and significance value are shown above each test with each data point representing an individual sample. (D) Tests reaching a significance of p<0.05 were further analyzed by ROC with the AUC and significance value shown. (E) T cell metrics predicting the same allo-HCT outcome were combined into a multi-parameter ROC analysis with the test, metrics utilized, and AUC values shown in the graphs. ROC = receiver-operator-characteristic; AUC = area under the curve; G0- G2 indicates highest grade of GVHD achieved by the patient.

From a total of 11 T cell metrics analyzed, we identified five metrics as having the capacity to predict relapse, GVHD (of any grade) or ≥ grade 2 GVHD. These included the number of CD45RO^+^ T cells and the frequencies of blasting CD45RO^+^ T cells, regulatory T cells (T_regs_), CD8^+^ T cells and DPTs (Fig 1B, Supplemental Fig 3). We first analyzed the samples collected between days 7 and 17 in three binary tests to determine if any biomarkers could differentiate between two allo-HCT outcomes (Fig 1C). Specific comparisons that were shown to be significant by t-test were further evaluated using a receiver-operator-characteristic (ROC) analysis (Fig 1D). We identified two predictive biomarkers of relapse (CD45RO T cells and frequency of Tregs), four predictive biomarkers of GVHD (CD45RO T cells and the frequency of Tregs, CD8 and DPTs) and two predictive biomarkers of ≥ grade 2 GVHD (frequency of Tregs and DPTs) (Fig 1C-D). Combining the individual T cell metrics into one multi-parameter analysis further increased their predictive potential as measured by the area-under-the-curve (AUC) values that reached 0.848, 0.736 and 0.701 for relapse, GVHD and ≥ grade 2 GVHD prediction respectively (Fig 1E).

A second analysis included patient samples collected between days 22 and 83 and were further normalized relative to each individual patient’s date of grade 1 or 2 GVHD diagnosis. For patients who did not develop GVHD, we normalized their data to day 54, which was the average time to GVHD diagnosis among our cohort. In this analysis, only three of the original five T cell biomarkers showed any predictive value with all three metrics predicting relapse (CD45RO T cells and the frequencies of CD8 and DPTs) and the increased frequency of CD8 and DPTs predicting both GVHD (of any grade) and ≥ grade 2 GVHD (Supplemental Fig 3). Combining the predictive metrics into one multi-parameter analysis as before further increased their predictive value with AUC values of 0.892, 0.741 and 0.734 for relapse, GVHD and ≥ grade 2 GVHD prediction respectively (Supplemental Fig 3).

#### Xenogeneic transplantation also reveals DPTs as a predictive biomarker of GVHD development

To replicate our clinical observational study, we utilized a human xenogeneic transplant model wherein we transplanted various human graft tissue into immunodeficient mice^31,59,60^. As we have published previously, xenogeneic GVHD (xeno-GVHD) is dose-dependent with each human graft source possessing their own LD^50^ dose and that the development of xeno-GVHD is not donor dependent^59^. Sampling the blood of mice transplanted with human PB-MNC at regular intervals, we divided mice into those that developed lethal xeno-GVHD and those that did not based on their survival at 12 weeks post-transplant (Fig 2A). We observed the presence of DPTs, which were not present in the starting healthy graft tissue, primarily developing in mice that would later develop lethal xeno-GVHD (Fig 2A). We confirmed that the development of DPTs is not dependent on a single human donor, are not artifacts of doublets by ImageStream analysis and that they develop even when isolated human T cells are used for transplantation (Fig 2A-B and Supplemental Fig 4). DPTs were present in a variety of xeno-GVHD target organs and developed as early as 1-week after transplant in each of the four graft sources we tested (Fig 2D-H). The increased frequency of DPTs was also a predictive biomarker of lethal xeno-GVHD development in this model system at 1-, 3- and 6-weeks post-transplant (Fig 2I). We further confirmed that the frequency of Tregs 1-week after transplant could act as a predictive biomarker of lethal xeno-GVHD but with the contribution of Tregs already well established in the GVHD literature, we chose to focus on the origin and function of DPTs for the remainder of this study (Supplemental Fig 5).

**Figure 2.**
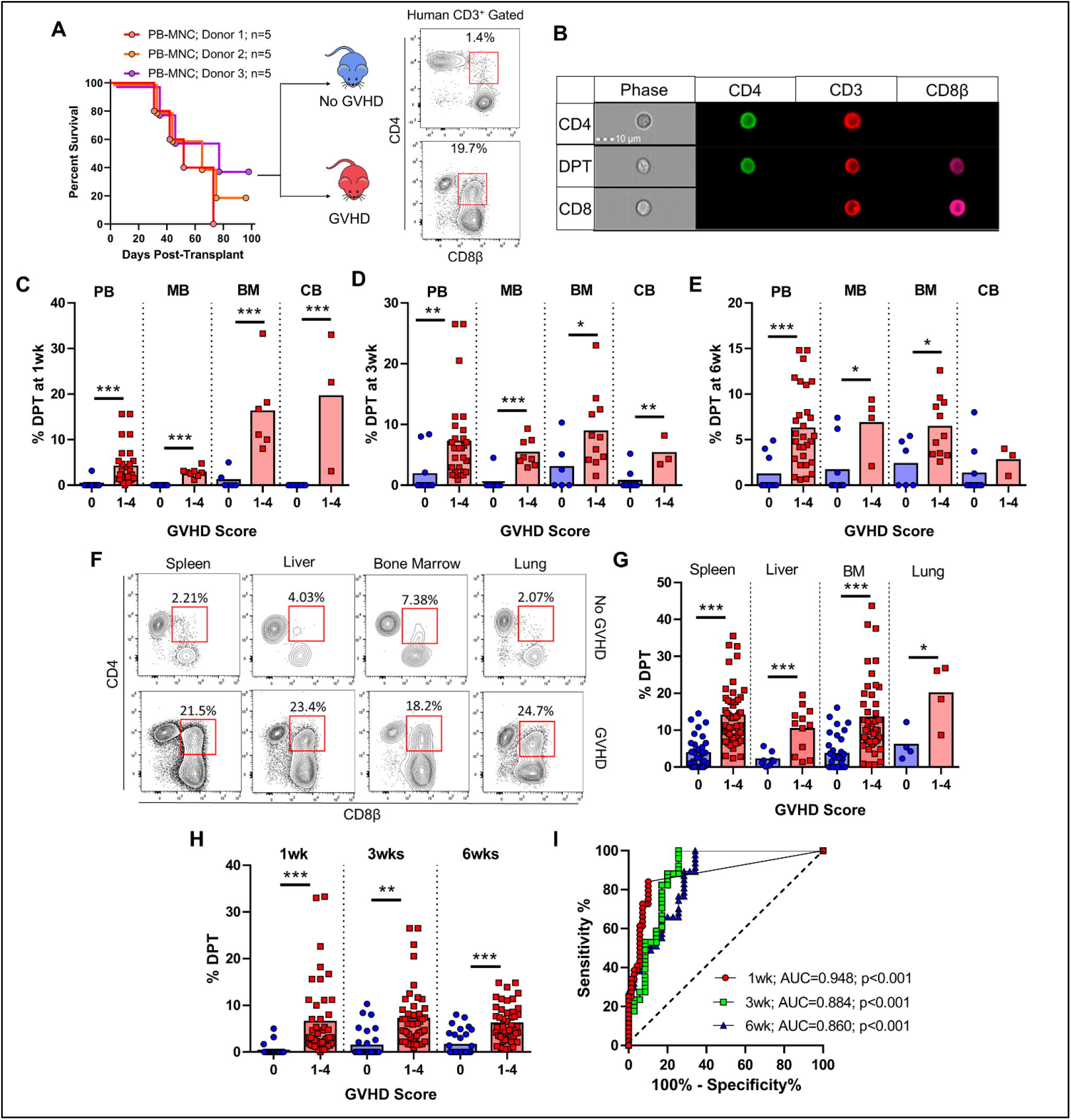
DPT develop after xenogeneic transplantation and are predictive of lethal GVHD. (A) Survival curve of mice transplanted with PB-MNC from three different donors. Mice were monitored for signs of xeno-GVHD and retrospectively assigned into the no GVHD (blue) or xeno-GVHD (red) group based on their survival at 12 weeks post-transplant. Representative dot plots are of human panHLA and CD3 gated mouse blood at 3 weeks post-transplant with the red box indicating the DPT gate. (B) Imagestream visualization of human T cells isolated from mouse blood. (C-E) Graphical representation of the percentage of DPTs in the blood of mice transplanted with the indicated graft source at 1wk (C), 3wks (D) and 6wks (E) post-transplant. (F-G) Dot plots and graph of the percentage of DPTs in the indicated organs of mice at the time of euthanasia. Red squares indicate DPT gate. (H-I) Blood data from (C-E) was combined (H) and analyzed by ROC (I) with the AUC and significance values indicated. Each point represents an individual mouse. PB (peripheral blood), MB (G-CSF mobilized peripheral blood), BM (bone marrow), CB (umbilical cord blood), MNC (mononuclear cells). Parametric t-test was used to determine significance. * p<0.05, ** p<0.01, *** p<0.001

#### The DPT population is transcriptionally distinct from CD4 and CD8 T cells

To explore if the CD4 and CD8 lineage markers are acquired via trogocytosis or de novo transcribed, we performed RNA-seq on flow-sorted CD4, CD8 and DPTs from xeno-GVHD mice with a sort-purity of >98% (Fig 3). Analysis of the transcriptome from these cells revealed that DPTs express a transcriptional signature unique from CD4 or CD8 T cells (Fig 3A-B). Using a threshold of p<0.01 and a fold change of >10 to determine significance, we reveal that DPTs differentially expressed 63 genes when compared to both CD4 and CD8 T cells (Fig 3A). Of the differentially expressed genes, we confirmed that both CD4 and CD8 are both transcriptionally expressed in DPTs (Fig 3C-E). The longevity of CD4, CD8α and CD8ß co-expression on isolated DPTs was also maintained in culture for up to three weeks and for at least 9 weeks when re-transplanted into naïve NSG mice. These data further suggest that co-expression of the CD4 and CD8 lineage markers is not an acquired feature but produced de novo (Supplemental Fig 6).

**Figure 3.**
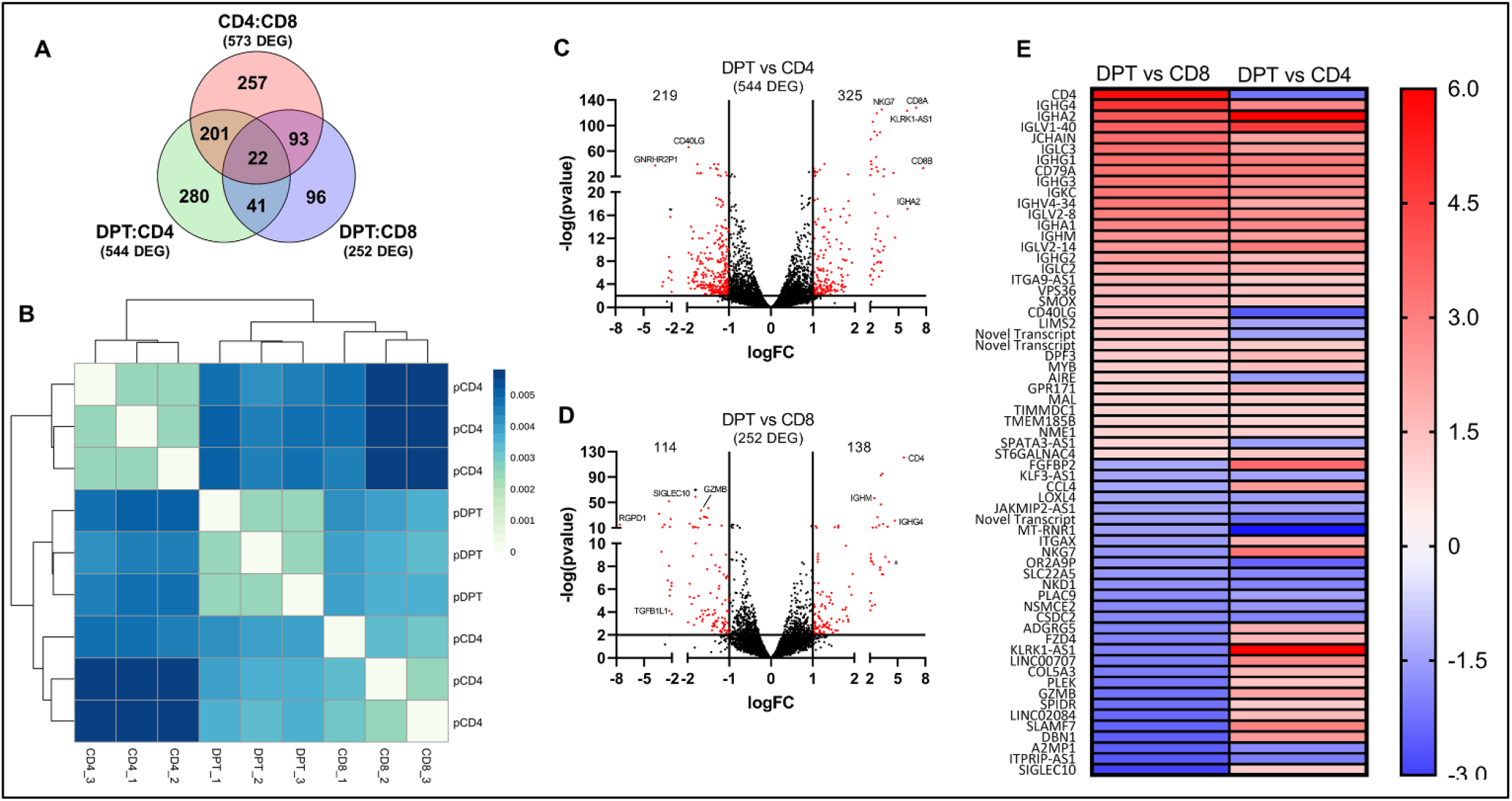
The transcriptome of DPT is unique relative to CD4 and CD8 T cells. (A) Venn diagram of the number of independent and shared differentially expressed genes among the three comparisons indicated. (B) Unsupervised hierarchical clustering of the RNA-seq samples using the computed Pearson correlation for each pair of samples and the Euclidean distances of the correlation distances determined. (C-D) Volcano plots showing the number of differentially expressed genes in the DPT:CD4 (C) and DPT:CD8 (D) comparisons using a cutoff of a ≥ 1 log expression difference and a p-value of 0.01. (E) List of the 63 genes differentially expressed in both the DPT:CD4 and DPT:CD8 comparisons with red indicating increased expression in DPTs relative to CD4/CD8 T cells and blue indicating decreased expression.

Using all 796 genes differentially expressed in the DPT:CD4 and DPT:CD8 comparisons, KEGG BRITE functional hierarchical classification highlighted most of the differentially expressed genes as having an enzymatic function, followed by CD molecules, exosome machinery, transcription factors and cytokine/cytokine receptors (Supplemental Table 2). Interestingly, DPTs also appear to express several genes normally associated with B cell biology, though the mechanism for this observation requires further investigation (Fig 3E). To confirm the differential expression of several inflammatory and inhibitory co-receptors, we surveyed 13 markers on the surface of CD4, CD8 and DPTs from xeno-GVHD mice (Fig 4A-C, Supplemental Fig 7). DPTs displayed an intermediate expression profile of CD27, OX40, PD-1 and LAG3 compared to CD4 and CD8 T cells analyzed from the blood or spleen of xeno-GVHD mice (Fig 4A-C, Supplemental Fig 6). Surprisingly, DPTs did not express elevated levels of the inhibitory markers TIGIT, TIM3 or CTLA4 commonly expressed on exhausted T cells. The expression of these markers was further explored on the T cells from our observational clinical study (Fig 4D-L). While the expression of these markers differed slightly, most likely due to differences in a murine versus human host environments, DPTs still displayed an expression profile of a terminally differentiated effector T cell population with little evidence of exhaustion (Fig 4D-L).

**Figure 4.**
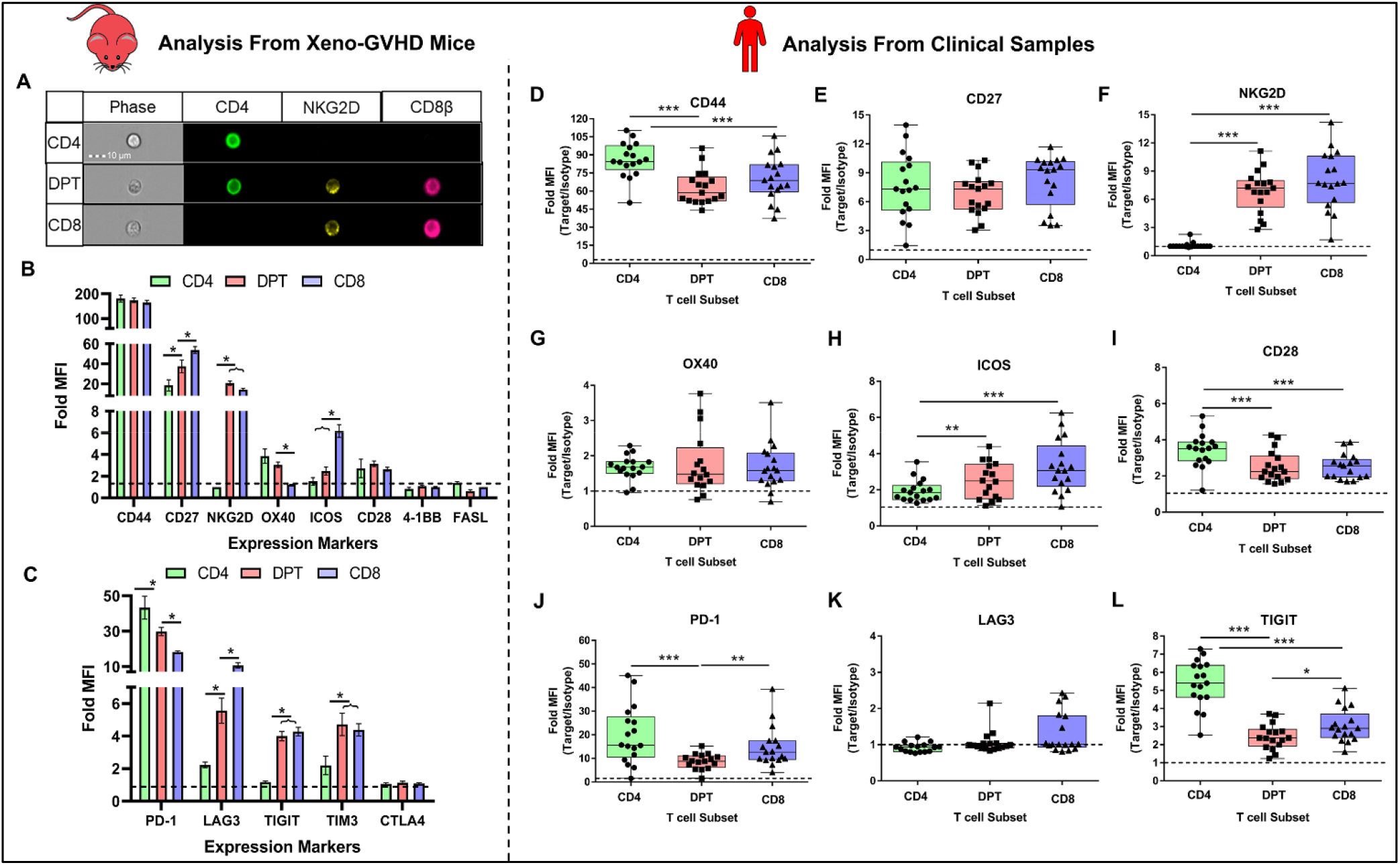
DPTs display a transitional profile with limited inhibitory receptor expression. (A) Imagestream visualization of DPTs expressing NKG2D. (B-C) Fold MFI of CD4 (green), DPTs (red) and CD8 T cells (blue) expression of eight pro-inflammatory (B) and five inhibitory (C) receptors analyzed from 5-8 xeno-GVHD mice at 3 weeks post-transplant. (D-L) Analysis of six pro-inflammatory (D-I) and three inhibitory receptors (J-L) on the surface of CD4 (green), DPT (red) or CD8 (blue) T cells from the allo-HCT patient samples collected as part of our observational study. Each dot represents the average Fold MFI of that marker across all samples collected for one individual patient. A minimum of 10 patients was collected for each marker. Expression of 4-1BB, FASL and CTLA4 had no expression over an isotype control and are not shown. TIM3 was not measured in the clinical samples. Error bars represent SEM. Fold MFI = median fluorescent intensity of marker / isotype control. Parametric t-test was used to determine significance. * p<0.05, ** p<0.01, *** p<0.001

#### Antigen stimulated CD8 T cells develop into DPTs

To identify the T cell population that gives rise to DPTs, we performed an unsupervised tSNE clustering of human T cells from a xeno-GVHD mouse using a nine-color flow cytometry panel (Fig 5A). This analysis, in addition to the global transcriptomic analysis, indicated that DPTs cluster more closely with CD8 T cells than CD4 T cells. Thus, despite the well documented plasticity of the CD4 lineage, our results suggest that DPTs may develop from the CD8 lineage (Fig 3B, Fig 5A).

**Figure 5.**
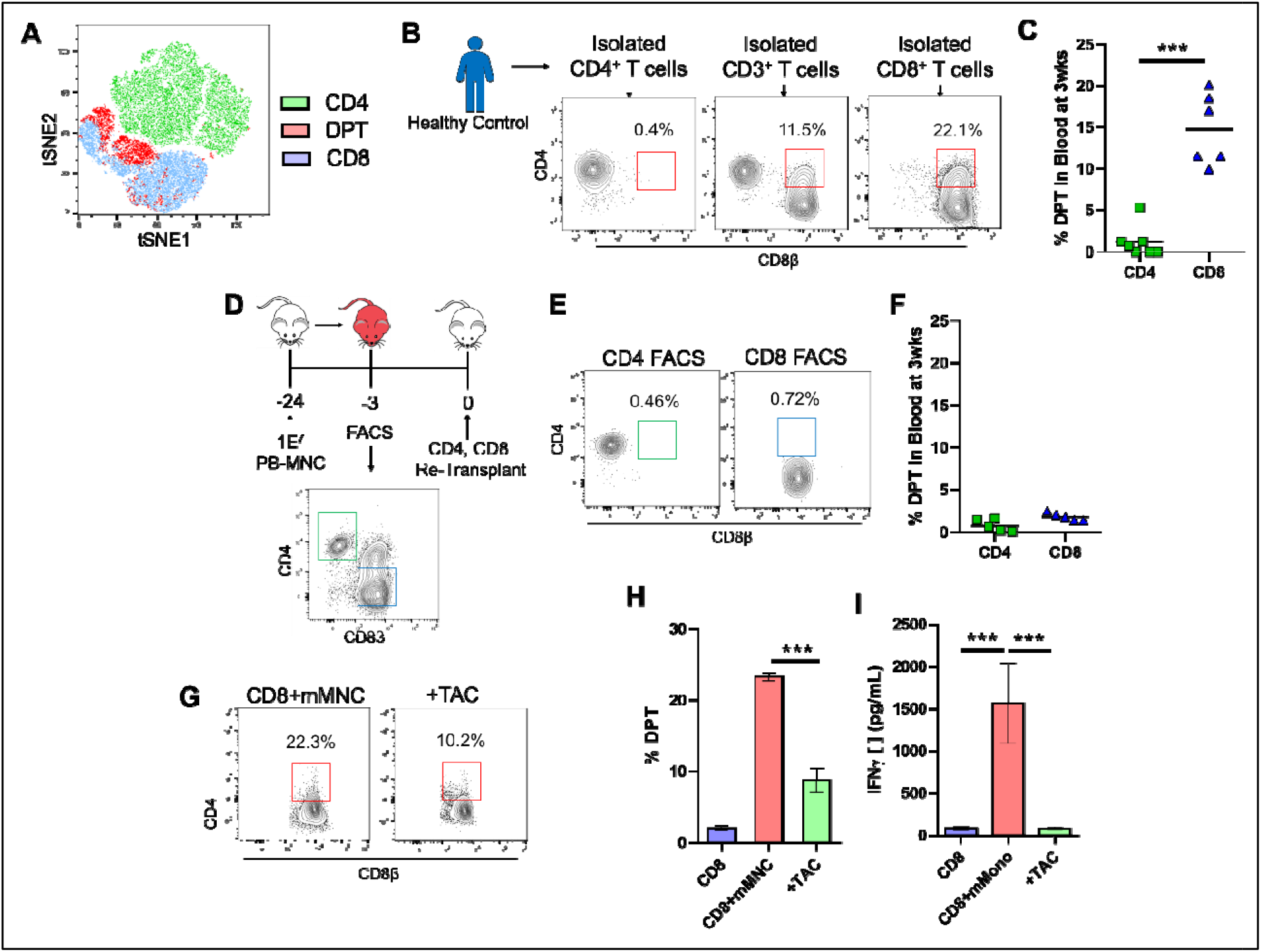
DPTs develop from CD8 T cells after antigenic stimulation. (A) Unsupervised tSNE clustering of human T cells concatenated from five xeno-GVHD mice. (B-C) Isolated CD3, CD4 and CD8 PB-Tc from a healthy donor were transplanted into mice. Dot plots show the percentage of DPTs in mouse blood at three weeks post-transplant (B) with the percentage of DPT quantified in (C). (D) Five mice transplanted with PB-MNC were euthanized and the human cells isolated prior to flow-sorting of CD4 and CD8 T cells. (E-F) Dot plots and bar graph of the percentage of DPTs in the blood of mice re-transplanted with flow-sorted CD4 and CD8 T cells. (G-I) CD8 T cells were co-cultured for five days with mouse MNC with or without tacrolimus and then analyzed for DPTs (H) and IFNγ (I). Parametric t-test was used to determine significance. * p<0.05, ** p<0.01, *** p<0.001

To test this hypothesis, we transplanted isolated CD4 or CD8 T cells from a healthy donor into our xenogeneic transplant model. Elevated frequencies of DPTs were only observed in mice transplanted with isolated CD8 T cells, indicating that mature CD8 T cells and not CD4 or progenitor T cells give rise to the DPT population (Fig 5B-C). To investigate if the development of DPTs is antigen-dependent or the result of a lymphopenic/xenogeneic environment, we first surveyed the phosphorylation status of several STAT proteins downstream of cytokine receptors commonly expressed by T cells. Unfortunately, we could not identify any differences in the phosphorylation of STAT1, 3, 4 and 6 between the CD8 and DPT populations. While the xenogeneic environment may still be playing a role in cytokine receptor/STAT activation, these data suggest that there is not any differential activation between the CD8 and DPT populations (Supplemental Fig 8A-B). To test the role of the xenogeneic environment further, we transplanted CD4 or CD8 T cells sorted from a xeno-GVHD mouse into a naïve mouse to determine if DPTs would reform after the secondary transplant. We could not detect any development of DPTs in the mice receiving sorted-CD8 T cells (Fig 5D-F). We next co-cultured CD8 T cells in the presence of mouse MNCs for five days with or without tacrolimus to inhibit TCR signaling. The data revealed that mouse MNCs in culture could induce both DPT development and IFNγ secretion which was inhibited by tacrolimus (Fig 5G-I). These data suggest that DPTs are developing as a result of antigenic/xenogeneic stimulation by the TCR and not as a direct result of a lymphopenic/xenogeneic environment.

Knowing that the transcriptome of DPTs is unique compared to CD8 T cells and that transcription factors were one of the functional classes highlighted in the KEGG BRITE analysis, we further investigated what transcription factors may be involved in the differentiation of DPT from the CD8 population. Unfortunately, we were not able to definitively identify a specific transcription factor that was correlated with the development of DPTs (Supplemental Fig 8C-G). Relative to CD4 T cells, the DPT population had higher transcriptional expression of RUNX3, T-bet and lower expression of FOXP3. Compared to CD8 T cells, only THPOK was statistically different with DPTs possessing a higher level of THPOK transcripts (Supplemental Fig 8C). To confirm these findings, we performed intracellular flow cytometry and western blotting on a subset of the transcriptional factors identified by the RNA-seq data. The protein level expression differences of RUNX3 were confirmed but were inconclusive regarding THPOK due to the overall low expression level of the transcription factor (Supplemental Fig 8E-G). Thus, while the transcriptional expression of THPOK in DPTs is interesting, additional studies will be needed to determine the importance of THPOK in DPT development.

#### Differentiated DPTs are a metabolically active and unique T cell population

To further explore DPT biology, we investigated their metabolic state relative to CD4 and CD8 T cells. We analyzed sorted CD4, CD8 and DPTs from xeno-GVHD mice using a two-photon (2P) autofluorescence imaging system (Fig 6A-E). This system is able to detect the autofluorescence of NAD(P)H and FAD at a single cell resolution and use their ratios to determine the optical redox ratio (ORR) of the cell population as a measurement of metabolic activity^61,62^. This analysis revealed that the DPT population had elevated ORR, indicative of high metabolic activity (Fig 6A-B). Furthermore, a random forest classifier trained to integrate the 10 metabolic parameters imaged, accurately identified the DPT population from CD4 and CD8 T cells shown as a confusion matrix, ROC curve and a UMAP dimensionality reduction (Fig 6C-E). Additional metabolic analysis using a Seahorse real-time metabolic analyzer independently confirmed that DPTs have a higher ATP production rate than CD4 and CD8 T cells with the increased ATP production primarily coming from glycolysis (Fig 6F). In addition to metabolic differences, we also analyzed the T cell populations for evidence of “blasting”. Blasting is characterized by increased cell size and is a marker of cellular proliferation in T cells. The DPT population displayed higher frequencies of the blasting phenotype relative to CD8 T cells in both xeno-GVHD and clinical GVHD samples (Supplemental Fig 9).

**Figure 6.**
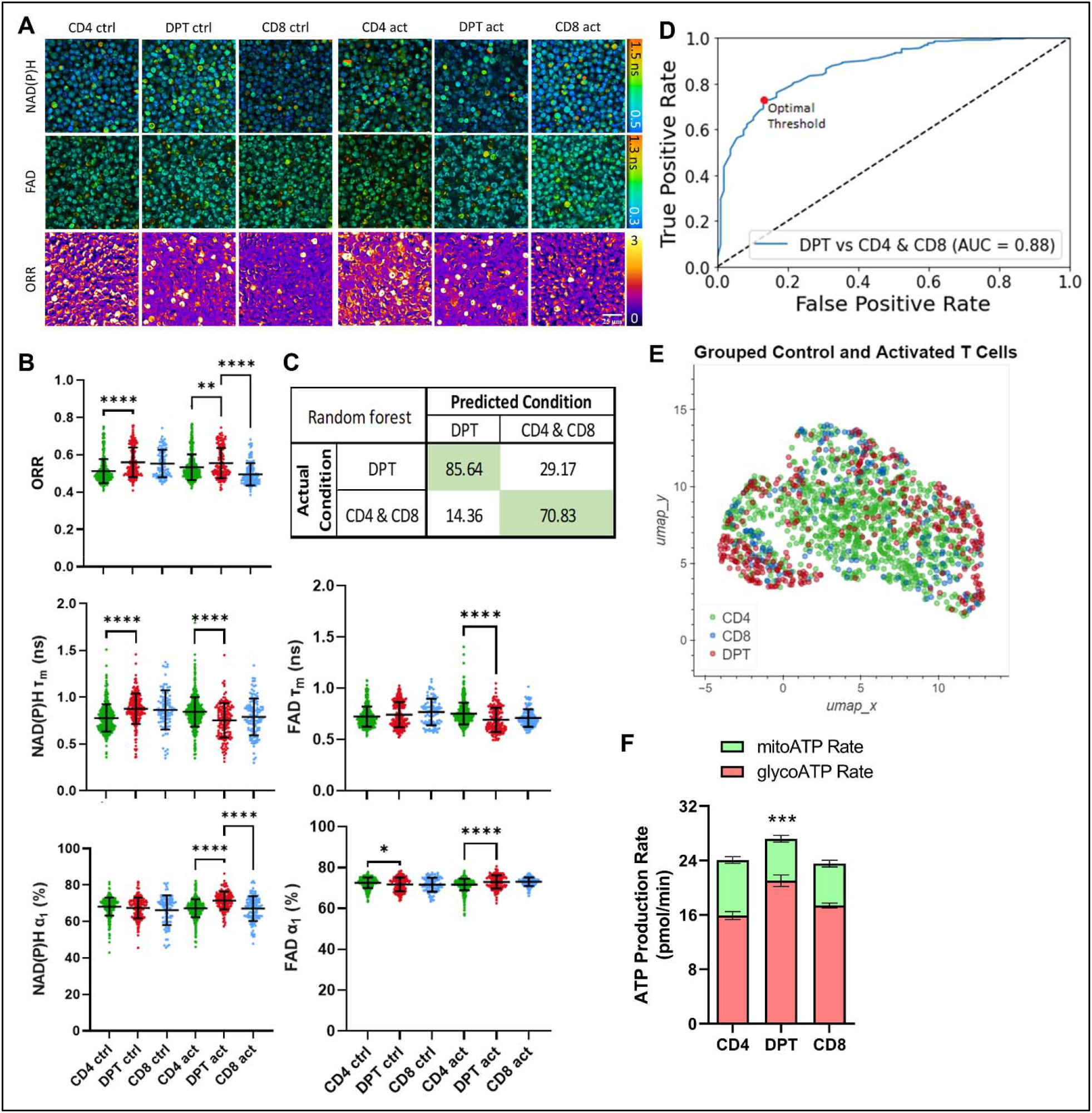
DPT cells exhibit a metabolically distinct signature from CD4 and CD8 T cells. (A) Representative images from live-cell imaging of NAD(P)H mean lifetime (τ_m_, 0.5-1.5 ns), FAD τ_m_ (0.3-1.3 ns), and optical redox ratio images from flow-sorted CD4, CD8 and DPT from xeno-GVHD mice. All cells were cultured for 48hrs prior to visualization with activated cells also stimulated with CD3/CD28/CD2 12hrs prior to imaging. Scale bar at top left, 25 μm. (B) Scatter plots of the optical redox ratio (NAD(P)H / (NAD(P)H + FAD), NAD(P)H τ_m_, FAD τ_m_, % unbound NAD(P)H, % protein-bound FAD. Each dot represents a single cell with mean at center and error bars representing 1 standard deviation. Confusion matrix (C) and ROC curve (D) of a random forest classifier trained on the 10 OMI parameters (NAD(P)H and FAD τ_1_, τ_2_, α_1_, α_2_, τ_m_). (E) UMAP dimensionality reduction analysis of OMI parameter results (τ_1_, τ_2_, α_2_, τ_m_) for both NAD(P)H and FAD. (F) Seahorse analysis of the ATP production rate of flow sorted CD4, DPT and CD8 T cells. Parametric t-test was used to determine significance. * p<0.05, ** p<0.01, *** p<0.001, **** p<0.0001. Cell counts ranged from 103-598 for each population.

#### DPTs represent a GVHD-specific T cell population with minimal GVL activity

To confirm the role of DPTs in the pathology of xeno-GVHD, we sorted CD4, CD8 and DPTs from xeno-GVHD mice and re-transplanted each population individually into naïve mice (Fig 7A). We observed that only the DPT population was sufficient to mediate lethal xeno-GVHD (Fig 7B-C). DPTs expanded rapidly after transplantation and had increased infiltration into the spleen, liver, and lungs relative to CD4 and CD8 T cells (Fig 7D-F). The transplanted DPT population primarily retained their DPT phenotype throughout the experiment (Supplemental Fig 6). Mice transplanted with DPTs also exhibited high levels of human IFNγ in their plasma compared to CD8 T cells (Fig 7G). To further investigate the secretome of DPTs, we cultured human T cells from xeno-GVHD mice overnight with brefeldin A and then stimulated with or without PMA/ionomycin prior to intracellular cytokine staining. DPTs secreted a diverse repertoire of cytokines including granzyme and perforin, the inflammatory modulators IL-17A, IL- 22 and GM-CSF as well as the activation cytokines IFNγ and TNFα, indicative of a highly inflammatory T cell population (Fig 7H-I, Supplemental Fig 10). To further support the role of DPTs in xeno-GVHD, we transplanted isolated CD8 T cells into NSG mice followed by two in vivo injections of a mouse anti-human CD4 antibody (200μg/injection) at days 5 and 10 post-transplant. We observed that using an anti-human CD4 antibody to deplete the DPT population largely protected mice from xeno-GVHD while all untreated mice developed lethal xeno-GVHD (Fig 7J). The anti-human CD4 antibody treatment effectively depleted the DPT population at 1- and 3-weeks post-transplant as shown by both flow cytometry of blood samples and the measurement of IFNγ in the plasma (Fig 7K-M). Antibody treatment did not alter the number of non-reactive CD8 T cells in the mouse (Fig 7N). The one mouse from the CD4 antibody treatment group that died at 9 weeks post-transplant had elevated levels of DPTs and IFNγ starting at 6 weeks, suggesting this mouse had only a partial depletion of the DPT population.

**Figure 7.**
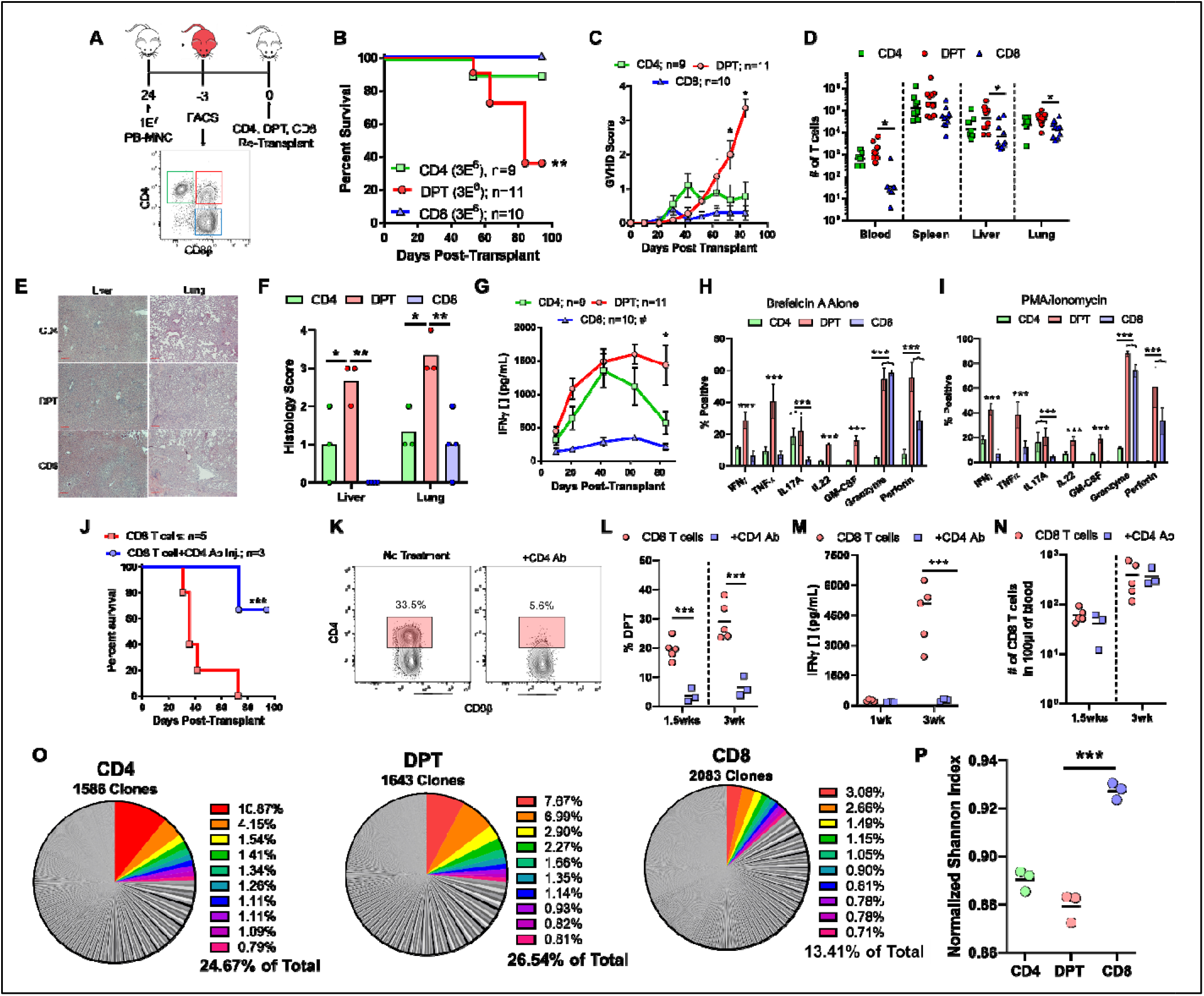
DPTs are sufficient to mediate xeno-GVHD pathology. Schematic of experimental design. Survival curve (B) and GVHD score (C) of mice transplanted with CD4, DPT and CD8 T cells previously isolated from five pooled xeno-GVHD mice. (D) Quantification of T cells in xeno-GVHD target organs at time of euthanasia. (E-F) Representative H&E staining of liver and lung sections from transplanted mice (E) and histological scoring (F). (G) Quantification of IFNγ levels in the plasma of transplanted mice. (HI) Splenic T cells from 4-13 xeno-GVHD mice were cultured overnight with either Brefeldin A alone (H) or Brefeldin A with a 4-hour PMA/ionomycin stimulation (I) prior to ICS staining of the indicated cytokine. (J-N) Analysis of mice transplanted with human CD8 T cells and treated with a mouse anti-human CD4 antibody (200μg/injection, retro-orbital) at days 5 and 10 posttransplant. Survival curve (J), dot plots (K) and graphical representation of the percentage of DPTs at 1.5- and 3-weeks (L). (M) Quantification of IFNγ in the plasma and the number of CD8 T cells in the blood at 1.5- and 3-weeks (N). (O-P) TCR clonal analysis of the TRB gene segment (J) and normalized Shannon-Weiner index calculation (K). Total number of unique TRB clones is indicated while the ten clones with the highest frequency are colored in the pie graph. Non-parametric t-tests were used for determining significance between T cell numbers while parametric t-tests were used for all other analyses. * p¼0.05, ** p¼0.01, *** p¼0.001

Clonal TCR analysis of the TRAB gene within the CD4, CD8 and DPT populations also provided evidence that the CD4 and DPT populations are experiencing clonal expansion. Both the CD4 and DPT populations had lower Shannon diversity indices and had the ten most abundant clones representing ~25% of the entire population (Fig 7O-P, Supplemental Table 3). While analysis of the TRAB locus supports the hypothesis that both CD4 and DPTs are experiencing clonal expansion and are being activated by a xenogeneic antigen, only the inflammatory DPT population was able to mediate lethal xeno-GVHD pathology.

While we have shown that DPTs are sufficient to mediate xeno-GVHD pathology, it is unclear if they can also have activity against human blood cancers. To investigate this question, we transplanted the human B-ALL cell line, NALM-6, into mice seven days prior to the transplantation of DPTs (sorted from xeno-GVHD mice) or freshly isolated human CD4 and CD8 T cells taken from the same donor (Fig 8A). The DPT population provided no survival advantage to the mice relative to mice not receiving any human T cells, while both the CD4 and CD8 T cell populations were able to delay death (Fig 8B). While all mice eventually succumbed to the cancer, mice receiving either CD4 or CD8 T cells had a lower cancer burden at three weeks post-transplant when compared to DPT or no treatment controls (Fig 8C-E). Failure of the DPT to provide protection was not the result of failed expansion or activation, as mice receiving DPTs had equivalent numbers of T cells as mice transplanted with CD4 or CD8 T cells and the presence of IFNγ in their plasma (Fig 8F-G). Interestingly, DPTs also exhibited a lower capacity for direct NALM-6 killing in an in vitro co-culture relative to CD4 and CD8 T cells (Fig 8H). To determine if this difference was due to DPTs having been sorted from a xeno-GVHD mouse, we repeated this experiment using CD4 and CD8 T cells sorted from xeno-GVHD mice. We observed the same phenotype, with DPTs failing to provide a survival advantage when challenged with either the human B-ALL cell line NALM-6 or RS4;11 (Supplemental Fig 11). Lastly, to determine if DPTs lacked GVL activity because they developed independent of human B-ALL cancer antigens, we repeated this experiment with isolated CD8 T cells and depleted the DPT population using a mouse anti-human CD4 antibody. Importantly, if DPTs also developed against human B-ALL, we should see a loss of protection when DPTs are depleted. As before, we did not observe any difference in GVL activity when the DPTs were depleted in regard to survival or the quantity of human B-ALL in the blood of mice (Fig 8I-J). Overall, these results suggest that this inflammatory DPT population, which develops from the CD8 T cell population after transplant as a result of antigen/xenogeneic stimulation by the TCR, may represent a T cell population that is biased toward causing GVHD pathology and possesses limited GVL activity.

**Figure 8.**
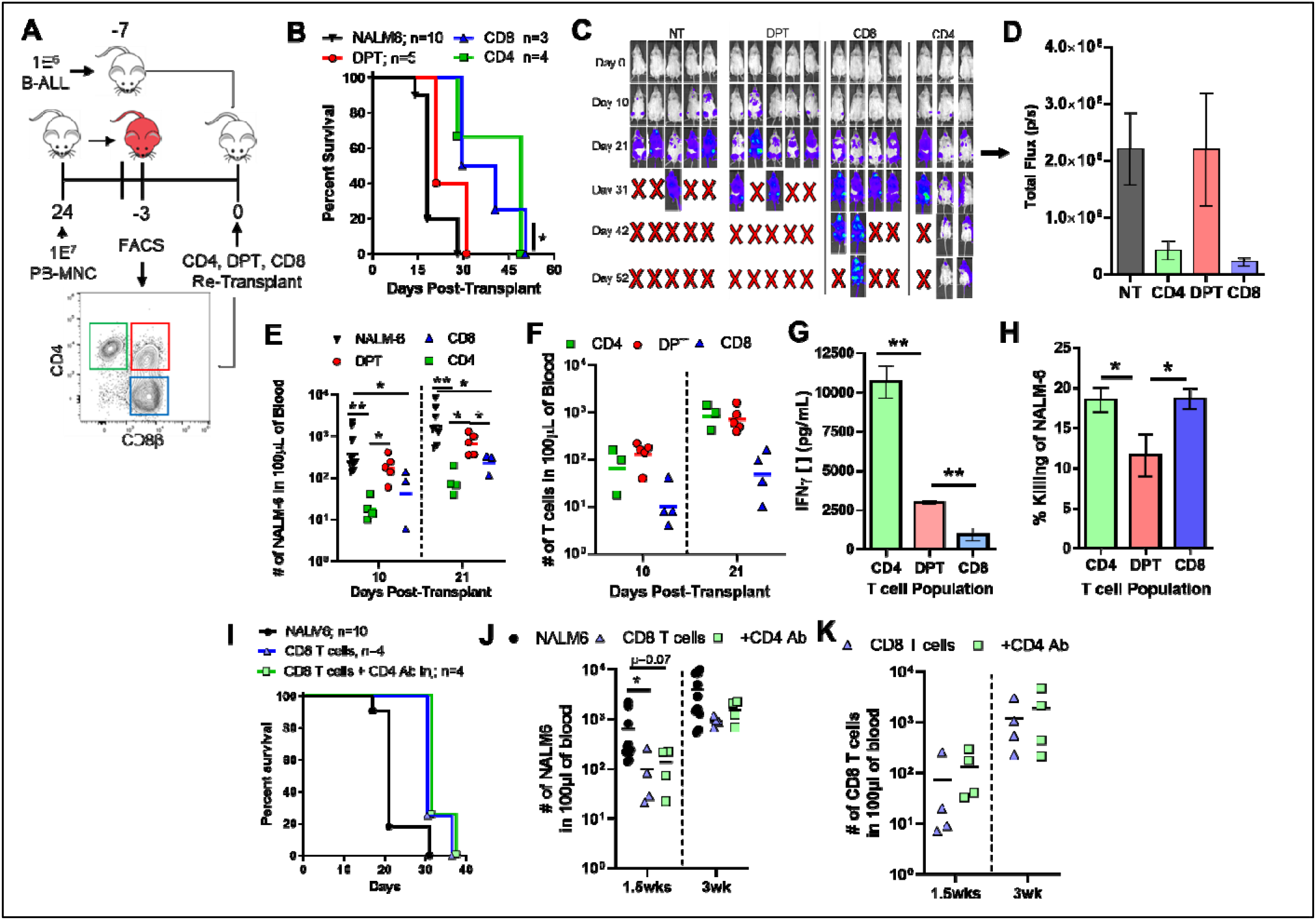
DPTs have limited GVL activity compared to CD4 and CD8 T cells. (A) Schematic of experiment design. The B-ALL cell line NALM-6 was transplanted into mice seven days prior to human T cells. DPTs were sorted from xeno-GVHD mice while fresh CD4and CD8 T cells were isolated from the same donor prior to transplantation. Survival curve (B),IVIS images (C), IVIS quantification (D), number of NALM-6 (E) and T cells (F) in the blood. IVIS quantification is for day 21 while NALM6 and T cell numbers at day 10 and 21 are shown. (G) Plasma IFNγ levels were also evaluated at three weeks post-transplant. (H) NALM-6 cells preloaded with Calcein dye were co-cultured with CD4, DPT and CD8 T cells for four hours and supernatants quantified for the presence of calcein. Data is expressed relative to a max kill condition using Triton-X. (I-K) Human CD8 T cells were transplanted followed by two retro-orbital injections of a mouse anti-human CD4 antibody (200μg/dose) at day 5 and 10 posttransplant. Survival curve (I), number of NALM6 cells in the blood of mice (J) and number ofCD8 T cells in the blood are shown (K). Non-parametric t-tests were used to determine significance for the quantification of B-ALL and T cell numbers while parametric t-tests were used for all other analyses. * p¼0.05, ** p¼0.01

### Discussion

In this study, we identified five T cell specific predictive biomarkers of allo-HCT outcome from a 35-patient observational study which included a unique CD4/CD8 double positive T cell (DPT) population as a predictor of GVHD. We further investigated this DPT population in a xenogeneic transplant model system and characterized the population as being transcriptionally, metabolically, and functionally distinct from CD4 and CD8 single-positive T cells. We also provide evidence that DPTs represent chronically activated CD8 T cells that gain the effector functions of the CD4 lineage. The highly inflammatory DPT population also showed sufficiency in mediating xeno-GVHD lethality but provided no observable survival advantage to mice challenged with human B-ALL malignancies. Thus, the human DPT population represents a highly activated and pathogenic T cell population directly implicated in the pathogenesis of GVHD but not GVL activity.

We are not the first group to identify the association of DPTs with xeno-GVHD. Alhaj Hussen et al also observed the development of DPTs during xeno-GVHD and that DPTs arise from the CD8 T cell population. While our study supports their findings that DPTs originate from the CD8 T cell population, their functional analysis led them to conclude that DPTs obtain a regulatory phenotype characterized by high levels of IL-10 and IL-13, low levels of IFNγ and high PD-1 expression^46^. Alternatively, our analysis of the DPT population suggestions that they are highly inflammatory, secreting IFNγ, TNFα, GM-CSF, IL-17A, IL-22, granzyme and perforin and are metabolically activated (Fig 6 and 7). Alhaj Hussen et al also found no association of DPTs with allo-HCT recipients with active GVHD when blood, duodenal or rectal biopsies were analyzed^46^. In our observational clinical study that analyzed blood samples from patients prior to GVHD diagnosis, we found that DPTs were strongly linked with future GVHD diagnosis (Fig 1).

It is not immediately clear why our two studies observed a different functional outcome for DPTs but there are several important differences to consider. Throughout our studies, we utilize CD8β as a marker of the CD8 lineage which is less promiscuous that CD8α^39^. For example, it has been shown by others that CD4 T cells can upregulate CD8α upon activation but not CD8²^39^. With their use of γ-irradiation prior to xenogeneic transplantation in Alhaj Hussen et al (a conditioning step we have shown to not be required for xeno-GVHD), it is possible that a proportion of the DPT population they identified actually represent activated CD4 T cells. This would explain why their DPT population had high levels of IL-10 (activated T_regs_) and IL-13 (TH_2_ response) while also having IFNγ and IL-17A expression (DPT response). Additionally, in our clinical cohort, we based our analysis on the CD45RO^+^ effector/memory T cell population that allowed us to remove the additional variability of non-reactive T cells. Re-analysis of our clinical data using the CD3^+^ T cell population (similar to Alhaj Hussen et al) resulted in lower resolution and weaker significance values across all of the biomarkers we tested, potentially explaining the difference between our clinical observations.

There are a variety of mechanisms that may explain the development of a human T cell population that co-expresses both CD8 and CD4 which include trogocytosis, recent thymic emigrants (RTEs), doublets and the transient expression of CD4 on activated CD8 T cells^58^. The use of a singlet gate during analysis and/or flow-sorting as well as Imagestream visualization exclude the possibility of DPTs being an artifact of improper gating (Fig 2C, Supplemental Fig 2). We have confirmed that DPTs arise irrespective of donor and graft source and the primary use of peripheral blood grafts in our study, which contain negligible numbers of RTEs and HSPCs, suggests DPTs are not RTE coming from the graft. Furthermore, NSG mice lack a functional thymus, excluding the possibility that DPTs are the result of de novo hematopoiesis^63^. Trogocytosis by CD8 T cells has been well documented but can also be excluded as DPTs arise after transplantation of isolated human CD8 T cells, wherein there is no source of human CD4 for them to acquire. The observation that CD4 is also upregulated transcriptionally suggests that the mechanism of CD4 acquisition is molecularly driven and not acquired from the environment.

Xenogeneic transplantation is a useful model to directly investigate the biology of human immune cells though we recognize that careful consideration must be given to xenogeneic transplantation studies. One important consideration is the nature of the antigen and antigen-presenting-cells (APC). While the specific antigen driving an allogeneic versus xenogeneic response is most likely different, the molecular response of the human T cell to the xeno-antigen can be informative on how human T cells respond to allo-antigen. As we have reviewed elsewhere, the interaction of the TCR/CD4/CD8 molecules with murine MHC is fully cross-reactive^31^. We have also shown in this study that the in vitro co-culture of human CD8 T cells with murine MNCs from NSG mice can drive the development of DPTs that can also be inhibited using tacrolimus (Fig 5G-I). Additionally, DPT development is not a direct consequence of the lymphopenic/xenogeneic environment as CD8 T cells isolated from a xeno-GVHD mice do not redevelop the DPT population after re-transplantation (Fig 5D-E). This, in addition to the decreased Shannon diversity index of the TCRs within the DPT versus CD8 populations, supports our conclusion that DPT development is driven by antigen stimulation (Fig 7J-K).

The T cell population is well known for its plasticity, with several other T cell populations identified in mice that share a similar function as DPTs. One population, denoted as Tc17 cells, are CD8 T cells that express a variety of CD4 lineage cytokines such as IL-17A, IL-22 and GM-CSF^45^. Furthermore, the directed depletion of this population was able to prevent lethal GVHD but were also shown to be non-cytolytic and have minimal GVL activity^45^. Two others studies describe how chronically activated CD4 T cells are able to co-express THPOK and RUNX3 with CD4 T cells differentiating into MHC-II restricted CD4+ cytotoxic effector cells^39,64^. While these T cell populations may be transcriptionally and/or functionally similar to the human DPTs we have explored in our study, there remains important differences between each of these populations which include the lack of CD4 expression in Tc17 cells and a difference in the starting population for CD4+ cytotoxic effector cells. One possibility is that these small but important differences are the result of subtle deviations between murine and human T cell biology with a directed study comparing murine Tc17 cells and human DPTs possibly required to fully understand their similarities/differences^65^.

In support of the xenogeneic experiments in this investigation, our 35-patient observational study of allo-HCT recipients also linked the presence of DPTs with GVHD diagnosis. Importantly, the 35 patients in our cohort all received post-transplant cyclophosphamide based GVHD prophylaxis, which is standard at our center. It is currently unclear if DPTs would also be predictive of GVHD development in allo-HCT patients receiving standard tacrolimus and methotrexate based GVHD prophylaxis. While patients were included in our cohort irrespective of HLA matching and conditioning regimen, additional studies will be required to validate DPTs as a predictive biomarker of GVHD in patients not treated with post-transplant cyclophosphamide (Supplemental Table 1).

To identify the five T cell predictive biomarkers in this study, we gated exclusively on the CD45RO^+^ T cell population (Supplemental Figure 2). The gating on CD45RO^+^ T cells is based on the knowledge that CD45RO is only expressed on effector and memory T cells and the hypothesis that gating on this effector/memory T cell population would provide additional sensitivity and specificity to the potential biomarkers. With the patient’s conditioning regimen eliminating or reducing their own T cell numbers to very low levels, the CD45RO^+^ T cell population most likely represents donor T cells from the graft. Thus, we are using CD45RO as a marker to identify donor effector T cells that are activated against either host or leukemia antigen. While we acknowledge that this gating strategy is not specific for effector T cells, it does allow us to remove naïve, non-reactive T cells from our analysis which improved the sensitivity of our biomarkers when compared to analysis of the entire T cell population.

Overall, this study supports the hypothesis that human DPTs are a highly inflammatory and pathogenic T cell population that are sufficient to mediate xeno-GVHD and strongly predictive of GVHD development in primary allo-HCT recipients. Metabolic, transcriptomic, and functional studies all identified DPTs as a T cell population with distinct inflammatory properties relative to single positive CD4 and CD8 T cells. This highly inflammatory state resulted in lethal xeno-GVHD pathology when isolated DPTs were transplanted into naïve mice while CD4 and CD8 T cells populations, also isolated from xeno-GVHD mice, failed to cause lethal pathology after re-transplantation. This inflammatory DPT population could not control two human B-ALL cell lines though, suggesting they have limited GVL activity. Furthermore, the increased clonality observed in DPTs compared to CD8 T cells and the capacity of tacrolimus to inhibit DPT development in vitro suggests that development of DPTs is the result of antigenic stimulation and not a direct result of a lymphopenic/xenogeneic host environment. Future studies will need to further explore the association of DPTs with other human chronic inflammatory disorders where they have already been identified including rheumatoid arthritis, hepatitis B and C, ²- thalassemia, renal carcinomas, breast cancers and islet graft rejection^47–53^. In conclusion, DPTs represent a unique T cell population whose presence should be directly investigated in other human chronic inflammatory diseases.

## Materials & Methods

### Study Design

The objective of this study was to identify T cell based predictive biomarkers of GVHD and relapse following allo-HCT and to describe the origin and function of human CD4+/CD8+ double positive T cells (DPTs). To achieve these aims, we performed an observational study of 35 allo-HCT recipients and performed several in vivo and in vitro experiments using a xenogeneic transplantation model system. We tested the hypothesis that DPTs arise from antigenic stimulation and that they are sufficient to mediate xenogeneic GVHD pathology. The number of donors, mice and replicates are indicated in the figure or figure legend. No data was excluded from this analysis. For mouse experiments, equal numbers of male and female mice were used in each experiment and specific experimental conditions were not grouped by cage to facilitate appropriate randomization.

### Isolation of Primary Human Cells

Human peripheral blood was collected from healthy consenting donors according to IRB protocol 2014-0806 (J.E.G.) and 2017-1070 (C.M.C). Leftover and de-identified remnants of human bone marrow and G-CSF mobilized peripheral blood grafts used for allo-HCT procedures at our clinic were collected under IRB protocol 2016-0298 (P.H.). De-identified human umbilical cord blood was collected from the Medical College of Wisconsin’s cord blood bank. All human blood samples were diluted 1:1 with leukocyte isolation buffer (PBS, 2mM EDTA, 2% FBS) prior to ficoll density-gradient centrifugation (1100xg for 15 min with 0 brake). When indicated, StemCell Technologies RosetteSep T cell, CD4 or CD8 enrichment kits were used.

### Clinical Sample Collection

All patients receiving an allo-HCT at the University of Wisconsin-Madison were prospectively enrolled between October 2020 and August 2021 on the minimal-risk IRB protocol 2020-1490 (C.M.C.). A total of 35 patients were enrolled in this study. Eligible patients had to receive post-transplant cyclophosphamide for GVHD prophylaxis at day +3/+4. Conditioning regimen, HLA matching status, graft source and patient disease type were all non-exclusion criteria for this study. Patients were divided into four respective categories based on their first reportable allo-HCT outcome: relapse of primary disease, no GVHD, grade 1 GVHD or grade 2-4 GVHD. Patient assignment into each respective GVHD category is based on their highest GVHD grade achieved between day 0 and 100 after transplant while a relapse event must have occurred within 1 year of transplant. In our cohort, one patient developed grade 2 GVHD at day 78, was treated with steroids and later relapse at day 176. Since it is unknown how the treatment of steroids influenced that patients rate of relapse, this patient was placed in the grade 2-4 GVHD group.

Excess blood from study patients was set aside after initial use and stored at 4 C for 3-10 days before processing and was never frozen. A total of 0.5mL of blood was aliquoted from each patient collection tube for each flow cytometry test and centrifuged at 600xg for 4min at 4 C. The cell pellet was resuspended in 1.5mL of 1X RBC lysis buffer (BioLegend), vortexed and rested for 2min prior to washing with 1mL of PBS and centrifugation. Supernatant was disposed of, and a second round of RBC lysis was performed. Afterwards, the cell pellet was re-suspended in 1.5mL of flow buffer (1X PBS + 2% FBS) and aliquoted into three flow cytometry tubes for staining.

### Transplantation of human cells into NSG Mice

All human cells were washed and resuspended in 150μL/mouse of sterile 1X PBS prior to retro-orbital injection. Equal numbers of male and female immunodeficient NSG (NOD.Cg-*Prkdc^scid^ Il2rg^tm1Wjl^/SzJ,* Jackson Laboratories) between the ages of 8-16 weeks of age were used for all experiments. All mice were weighed and monitored weekly for visible signs of GVHD with a scoring system carried out as follows: 0 = no signs of GVHD; 1 = 2-5% weight loss; 2 = 6-9% weight loss; 3 = 10-14% weight loss; 4 = ≥ 15% weight loss. Blood was regularly drawn at 1, 3, 6, 9- and 12-weeks post-transplant by retro-orbital bleeding. At time of euthanasia, the spleen, one lobe of the liver, lungs and femur were regularly collected for further processing.

### Flow Cytometry and Cell-Sorting

All cells were stained in flow buffer (PBS, 10% FBS) prior to quantification on a ThermoFisher Attune NxT Flow Cytometer and analysis on FlowJo V10. Antibody clones include mCD45 (A20), ICOS (C398.4A), T-bet (4B10), LAG3 (11C3C65), FASL (NOK-1), CD4 (OKT4), CD69 (FN50), OX40 (Ber-ACT35), CD27 (O323), pSTAT4 (p38), RORyt (Q21-559), CD44 (BJ18), NKG2D (1D11), PD-1 (EH12.2H7), THPOK (ZFP-67), TIGIT (A15153G), CD45RO (UCHL1), TIM3 (F38-2E2), pSTAT6 (A15137E), IL-22 (2G12A41), GATA3 (16E10A23), CD8α (RPA-T8), CD8² (SIDI8BEE), CD25 (BC96), 4-1BB (4B4-1), CD28 (CD28.2), RUNX3 (SD0803), CTLA4 (BNI3), PanHLA (w6/32), CD3 (UCHT1). Visualization of cells was complete using an ImageStream Mark II.

For cell sorting, cells were collected from the blood, spleen, liver and lungs of mice transplanted with a lethal dose of PB-MNC approximately 4 weeks prior. Processed tissues were ficolled as described above to obtain mononuclear cells. Human cells were isolated from the combined mononuclear cells using StemCell Technologies EasySep Mouse/Human Chimera isolation kit. Human cells were then stained in sterile PBS with the human CD4 (OKT4) and CD8² (SIDI8BEE) antibodies prior to sorting on a BD FACSAria Cell Sorter. Cells were then maintained in cell culture for 2-3 days prior to transplantation and/or expansion.

### Cell Lines and Cell Culture

The human B-ALL line RS4;11 and NALM-6 were purchased from ATCC and cultured using standard RPMI-10 media (RPMI base media, 10% FBS, 1X Pen/Strep, 1mM NEAA, 1X GlutaMAX). After flow sorting, isolated human CD4, CD8 and DPT populations were expanded by co-culturing with a 10X dose of γ-irradiated (25Gy) human mononuclear cells from a third-party donor with 5μg/mL of PHA in RPMI-10 media supplemented with 5ng/mL of human IL-2 and IL-7. RPMI-10 media supplemented with IL-2 and IL-7 was added/changed approximately every 3-4 days. Human CD8 T cells were co-cultured with mouse MNC isolated from the spleen, liver and lungs of NSG mice after ficoll separation. Cells were cultured with IL-2 and IL-7 as previously highlighted. Tacrolimus was added at a concentration of 20μM (MedChemExpress).

### RNA sequencing

T cells from five GVHD mice were pooled prior to flow-sorting of the CD4, CD8 and DPTs. Pooled T cell populations were then split into three replicates for RNA isolation. All RNA isolation and sequencing was performed by the Gene Expression Center at the University of Wisconsin-Madison. RNA was isolated with Qiagen RNeasy kit prior to a quality control check using an Agilent Bioanalyzer and NanoDrop One spectrophotometer. mRNA libraries were prepared using the TruSeq stranded mRNA kit and sequenced on an Illumina NovaSeq 6000 platform to 30 million reads per sample. Sample reads were processed by the University of Wisconsin Biotechnology Center.

### ELISAs

Mouse blood samples were diluted with 50μL of 1X PBS prior to centrifugation with 60μL of the diluted plasma stored at −20°C. Plasma was further diluted 1:5 prior to ELISA with Assay Buffer (PBS, 1mM Tween-20, 5% FBS). IFNγ ELISAs used the MD-1 clone for coating and a biotinylated 4S.B3 clone for detection prior to visualization with a streptavidin-HRP antibody (BioLegend).

### Western Blot

Equivalent number of T cells (1E6 cells/group) were lysed in Laemmli Sample Buffer with ²-mercaptoethanol (Bio-Rad, CA). Total cell lysate for each sample were resolved on Bis-Tris 4–12% gel (ThermoFisher NuPAGE) and transferred to polyvinylidene fluoride membranes (Millipore, Billerica, MA). The membranes were blocked in LI-COR blocking buffer (LI-COR, NE), Immunoblotting was performed by incubating the membranes with anti-human RUNX3 (Mouse, Cell Signaling Tech, MA), anti-human ThPOK (Rabbit, Cell Signaling Tech, MA), and anti-human GAPDH (Rabbit, Cell Signaling Tech, MA), according to the manufacturer’s recommendations. The membranes were then washed with TBST and incubated with fluorescent secondary antibodies (LI-COR, NE) and the immunoreactive bands were visualized using the Odyssey ^®^ CLx imaging system (LI-COR, NE).

### Two-photon lifetime imaging of NAD(P)H and FAD

A custom-built inverted multiphoton microscope (Bruker Fluorescence Microscopy, Middleton, WI, USA), was used to acquire fluorescence intensity and lifetime images. The equipment consists of an ultrafast laser (Spectra Physics, Insight DSDual), an inverted microscope (Nikon, Eclipse Ti), and a 40X water immersion (1.15 NA, Nikon) objective. NAD(P)H and FAD images were obtained for the same field of view. Using the masks, single cell features were extracted including bi-exponential fit parameters (τ_1_, τ_2_, α_1_, α_2_) for NAD(P)H and FAD, NAD(P)H τm, FAD τ_m_, and the optical redox ratio. Features were calculated on a per-cell level by averaging all the pixels across each cell. A minimum of 100 cells were analyzed per condition to obtain statistically robust results.

Additional information can be found in the Supplemental Material.

### Seahorse Metabolic Assay

ATP production rate was calculated from flow sorted CD4, CD8 and DPT using an Agilent Seahorse XF Real-Time ATP Rate Assay Kit using a Seahorse XF/XFe96 analyzer and following the manufacturer’s instructions.

### Bioluminescent Imaging

A total of 1E^6^ luciferase positive NALM-6 cells were injected retro-orbitally seven days prior to human T cell transplantation as described above. Luminescence detection was gathered at the indicated times using an IVIS Spectrum In Vivo Imaging System (PerkinElmer). Briefly, mice were anesthetized and intraperitoneally injected with 100μL of D-luciferin at 30mg/mL with luminescence obtained approximately 15 minutes later.

### Statistics

Graphs and statistical tests were completed using GraphPad Prism 6. All significance tests were performed using either unpaired parametric (for linear data) or non-parametric (for logarithmic data) t-tests. For the biomarker multivariate analysis, we used the multiplications of the empirical cumulative distribution factor (ecdf) for each variable to achieve a combined score with the R language. The AUC-ROC was used to determine the optimal direction of the exdf (< or >) for each variable. Generalization performance was estimated using leave-one-out crossvalidation. For the RNA-seq analysis each fastq file is assessed for quality control issues and trimmed to remove adapter sequences that were not removed by the initial demultiplexing. After trimming and QC, the reads were aligned to the human genome using the splice-junction aware read aligner STAR. The software package RSEM was used to generate normalized read counts for each gene and its potential isoforms. Genes with very low expression, which would otherwise reduce statistical power, were filtered out. The program EdgeR is then employed to perform differential gene expression analysis.

### Study Approval

All work involving human cells and tissues was performed in accordance with IRB protocol 2014-0806 (C.M.C.). All animal work was performed in accordance with UW-Madison IACUC protocol M005915 (C.M.C.).

## Supplementary Materials

This pdf includes Figures S1 to S11, Tables S1 to S3 and additional methodology on the collection of NAD(P)H and FAD autofluorescence using two-photon lifetime imaging.

## Acknowledgements

The authors would like to thank the Clinical Laboratory team at UW-Health including Hugo Stoffel-Rosales, Robin Rustad and Kaitlyn Walker for their help with sample collection, as well as the the University of Wisconsin Carbone Cancer Center (UWCCC) Flow Cytometry Core Facility, UWCCC Small Molecule Screening Facility, UWCCC Small Animal Imaging and Radiotherapy Facility, UWCCC Experimental Animal Pathology Laboratory and UWCCC Informatics Shared Resource, who are all supported by NIH/NCI P30-CA014520. The authors also utilized the University of Wisconsin-Madison Biotechnology Center Bioinformatics Core Facility for the analysis of RNA-sequencing data.

## Author Contributions

NJH, JEG, PH and CMC conceptualized the study; NJH, DT, JR, SJM and ECG performed and analyzed experiments; KN, AWH, NC, MCS JEG and PH provided supervision; NJH and CMC drafted, reviewed, and edited the manuscript. All authors revised and approved the manuscript.

## Data and Materials Availability

All data needed to evaluate the conclusions in this paper are present in the paper and/or the Supplemental Materials.

## References

1. D’Souza A, Fretham C, Lee SJ, et al. Current Use of and Trends in Hematopoietic Cell Transplantation in the United States. Biol Blood Marrow Transplant. 2020;26(8):e177–e182.

2. Gooptu M, Romee R,St Martin A, et al. HLA-haploidentical vs matched unrelated donor transplants with posttransplant cyclophosphamide-based prophylaxis. Blood. 2021;138(3):273–282.

3. Kanate AS, Majhail NS, Savani BN, et al. Indications for Hematopoietic Cell Transplantation and Immune Effector Cell Therapy: Guidelines from the American Society for Transplantation and Cellular Therapy. Biol Blood Marrow Transplant. 2020;26(7):1247–1256.

4. Martinez-Cibrian N, Zeiser R, Perez-Simon JA. Graft-versus-host disease prophylaxis: Pathophysiology-based review on current approaches and future directions. Blood Reviews. 2020;100792.

5. Ciurea SO, Zhang M-J, Bacigalupo AA, et al. Haploidentical transplant with posttransplant cyclophosphamide vs matched unrelated donor transplant for acute myeloid leukemia. Blood. 2015;126(8):1033–1040.

6. Battipaglia G, Labopin M, Kröger N, et al. Posttransplant cyclophosphamide vs antithymocyte globulin in HLA-mismatched unrelated donor transplantation. Blood. 2019;134(11):892–899.

7. Blazar BR, Hill GR, Murphy WJ. Dissecting the biology of allogeneic HSCT to enhance the GvT effect whilst minimizing GvHD. Nat Rev Clin Oncol. 2020;

8. Chang Y-J, Zhao X-Y, Huang X-J. Strategies for Enhancing and Preserving Anti-leukemia Effects Without Aggravating Graft-Versus-Host Disease. Front Immunol. 2018;9:3041.

9. Reddy P, Maeda Y, Liu C, et al. A crucial role for antigen-presenting cells and alloantigen expression in graft-versus-leukemia responses. Nat. Med. 2005;11(11):1244–1249.

10. Meurer T, Crivello P, Metzing M, et al. Permissive HLA-DPB1 mismatches in HCT depend on immunopeptidome divergence and editing by HLA-DM. Blood. 2021;137(7):923–928.

11. Mayor NP, Wang T, Lee SJ, et al. Impact of Previously Unrecognized HLA Mismatches Using Ultrahigh Resolution Typing in Unrelated Donor Hematopoietic Cell Transplantation. JCO. 2021;JCO.20.03643.

12. Petersdorf EW, Stevenson P, Bengtsson M, et al. HLA-B leader and survivorship after HLA-mismatched unrelated donor transplantation. Blood. 2020;136(3):362–369.

13. Petersdorf EW, Gooley TA, Malkki M, et al. HLA-C expression levels define permissible mismatches in hematopoietic cell transplantation. Blood. 2014;124(26):3996–4003.

14. Fuchs EJ, McCurdy SR, Solomon SR, et al. HLA Informs Risk Predictions after Haploidentical Stem Cell Transplantation with Post-transplantation Cyclophosphamide. Blood. 2021;blood.2021013443.

15. Wachsmuth LP, Patterson MT, Eckhaus MA, et al. Post-transplantation cyclophosphamide prevents graft-versus-host disease by inducing alloreactive T cell dysfunction and suppression. J Clin Invest. 2019;129(6):2357–2373.

16. Nunes NS, Kanakry CG. Mechanisms of Graft-versus-Host Disease Prevention by Posttransplantation Cyclophosphamide: An Evolving Understanding. Front Immunol. 2019;10:2668.

17. Paczesny S. Biomarkers for posttransplantation outcomes. Blood. 2018;131(2ũ):2193–2204.

18. Chen S, Zeiser R. Novel Biomarkers for Outcome After Allogeneic Hematopoietic Stem Cell Transplantation. Front Immunol. 2020;11:1854.

19. Adom D, Rowan C, Adeniyan T, Yang J, Paczesny S. Biomarkers for Allogeneic HCT Outcomes. Front Immunol. 2020;11:673.

20. Srinagesh HK, Özbek U, Kapoor U, et al. The MAGIC algorithm probability is a validated response biomarker of treatment of acute graft-versus-host disease. Blood Adv. 2019;3(23):4034–4042.

21. Hotta M, Satake A, Yoshimura H, et al. Elevation of Early Plasma Biomarkers in Patients with Clinical Risk Factors Predicts Increased Nonrelapse Mortality after Allogeneic Hematopoietic Stem Cell Transplantation. Transplant Cell Ther. 2021;27(8):660.e1–660.e8.

22. Rowan CM, Pike F, Cooke KR, et al. Assessment of ST2 for risk of death following graft-versus-host disease in pediatric and adult age groups. Blood. 2020;135(17):1428–1437.

23. Major-Monfried H, Renteria AS, Pawarode A, et al. MAGIC biomarkers predict long-term outcomes for steroid-resistant acute GVHD. Blood. 2018;131(25):2846–2855.

24. Zewde MG, Morales G, Gandhi I, et al. Evaluation of Elafin as a Prognostic Biomarker in Acute Graft-versus-Host Disease. Transplantation and Cellular Therapy, Official Publication of the American Society for Transplantation and Cellular Therapy. 2021;27(12):988.e1–988.e7.

25. Subburaj D, Ng B, Kariminia A, et al. Metabolomic identification of α-ketoglutaric acid elevation in pediatric chronic graft-versus-host disease. Blood. 2022;139(2):287–299.

26. Inamoto Y, Martin PJ, Lee SJ, et al. Dickkopf-related protein 3 is a novel biomarker for chronic GVHD after allogeneic hematopoietic cell transplantation. Blood Adv. 2020;4(11):2409–2417.

27. McCurdy SR, Radojcic V, Tsai H-L, et al. Signatures of GVHD and Relapse after Post-Transplant Cyclophosphamide Revealed by Immune Profiling and Machine Learning. Blood. 2021;blood.2021013054.

28. Cruz CRY, Bo N, Bakoyannis G, et al. Antigen-specific T cell responses correlate with decreased occurrence of acute GVHD in a multicenter contemporary cohort. Bone Marrow Transplant. 2021;

29. Podgorny PJ, Liu Y, Dharmani-Khan P, et al. Immune cell subset counts associated with graft-versus-host disease. Biol Blood Marrow Transplant. 2014;20(4):450–462.

30. Magenau JM, Qin X, Tawara I, et al. Frequency of CD4+CD25hiFOXP3+ Regulatory T Cells Has Diagnostic and Prognostic Value as a Biomarker for Acute Graft-versus-Host-Disease. Biology of Blood and Marrow Transplantation. 2010;16(7):907–914.

31. Hess NJ, Brown ME, Capitini CM. GVHD Pathogenesis, Prevention and Treatment: Lessons From Humanized Mouse Transplant Models. Frontiers in Immunology. 2021;12:3082.

32. Hill GR, Betts BC, Tkachev V, Kean LS, Blazar BR. Current Concepts and Advances in Graft-Versus-Host Disease Immunology. Annu Rev Immunol. 2021;

33. TeshimaT, Hill GR. The Pathophysiology and Treatment of Graft-Versus-Host Disease: Lessons Learnt From Animal Models. Front Immunol. 2021;12:715424.

34. Piper C, Zhou V, Komorowski R, et al. Pathogenic Bhlhe40÷ GM-CSF+ CD4+ T Cells Promote Indirect Alloantigen Presentation in the GI Tract during GVHD. Blood. 2019;

35. Gartlan KH, Bommiasamy H, Paz K, et al. A critical role for donor-derived IL-22 in cutaneous chronic GVHD. Am. J. Transplant. 2018;lδ(4):810–820.

36. Piper C, Hainstock E, Yin-Yuan C, et al. Single cell immune profiling reveals a developmentally distinct CD4+ GM-CSF+ T cell lineage that induces GI tract GVHD. Blood Adv. 2022;bloodadvances.2021006084.

37. Park S, Griesenauer B, Jiang H, et al. Granzyme A-producing T helper cells are critical for acute graft-versus-host disease. JCI Insight. 2020;

38. Hashimoto K, Kouno T, Ikawa T, et al. Single-cell transcriptomics reveals expansion of cytotoxic CD4 T cells in supercentenarians. Proc Natl Acad Sci USA. 2019;116(48):24242–24251.

39. Mucida D, Husain MM, Muroi S, et al. Transcriptional reprogramming of mature CD4^+^ helper T cells generates distinct MHC class II-restricted cytotoxic T lymphocytes. Nat. Immunol. 2013;14(3):281—289.

40. Takeuchi A, Saito T. CD4 CTL, a Cytotoxic Subset of CD4+ T Cells, Their Differentiation and Function. Front Immunol. 2017;8:.

41. Dwyer GK, Mathews LR, Villegas JA, et al. IL-33 acts as a costimulatory signal to generate alloreactive Thl cells in graft-versus-host disease. J Clin Invest. 2022;e150927.

42. Carlson MJ, West ML, Coghill JM, et al. In vitro-differentiated TH17 cells mediate lethal acute graft-versus-host disease with severe cutaneous and pulmonary pathologic manifestations. Blood. 2009;113(6):1365–1374.

43. Agle K, Vincent BG, Piper C, et al. Bim regulates the survival and suppressive capability of CD8+ FOXP3+ regulatory T cells during murine GVHD. Blood. 2018;132(4):435–447.

44. Beres AJ, Haribhai D, Chadwick AC, et al. CD8+ Foxp3÷ regulatory T cells are induced during graft-versus-host disease and mitigate disease severity. J. Immunol. 2012;189(1):464–474.

45. Gartlan KH, Markey KA, Varelias A, et al. Tc17 cells are a proinflammatory, plastic lineage of pathogenic CD8+ T cells that induce GVHD without antileukemic effects. Blood. 2015;126(13):1609–1620.

46. Alhaj Hussen K, Michonneau D, Biajoux V, et al. CD4+CD8+ T-Lymphocytes in Xenogeneic and Human Graft-versus-Host Disease. Front Immunol. 2020;11:579776.

47. Menard LC, Fischer P, Kakrecha B, et al. Renal Cell Carcinoma (RCC) Tumors Display Large Expansion of Double Positive (DP) CD4+CD8+T Cells With Expression of Exhaustion Markers. Front Immunol. 2018;9:2728.

48. Choi YJ, Park H-J, Park HJ, Jung KC, Lee J-I. CD4hiCD81ow Double-Positive T Cells Are Associated with Graft Rejection in a Nonhuman Primate Model of Islet Transplantation. J Immunol Res. 2018;2018:3861079.

49. Zahran AM, Saad K, Elsayh KI, Alblihed MA. Characterization of circulating CD4+ CD8+ double positive and CD4- CD8- double negative T-lymphocyte in children with β-thalassemia major. Int. J. Hematol. 2017;105(3):265–271.

50. Quandt D, Rothe K, Scholz R, Baerwald CW, Wagner U. Peripheral CD4CD8 double positive T cells with a distinct helper cytokine profile are increased in rheumatoid arthritis. PLoS ONE. 2014;9(3):e93293.

51. Nascimbeni M, Pol S, Saunier B. Distinct CD4+CD8+ Double-Positive T Cells in the Blood and Liver of Patients during Chronic Hepatitis B and C. PLOS ONE. 2011;6(5):e20145.

52. Desfrançois J, Derré L, Corvaisier M, et al. Increased frequency of nonconventional double positive CD4CD8 αβT cells in human breast pleural effusions. International Journal of Cancer. 2009;125(2):374–380.

53. Nascimbeni M, Shin E-C, Chiriboga L, Kleiner DE, Rehermann B. Peripheral CD4+CD8+ T cells are differentiated effector memory cells with antiviral functions. Blood. 2004;104(2):478–486.

54. Sullivan YB, Landay AL, Zack JA, Kitchen SG, Al-Harthi L. Upregulation of CD4 on CD8+ T cells: CD4dimCD8bright T cells constitute an activated phenotype of CD8+ T cells. Immunology. 2001;103(3):270–280.

55. Virdi AK, Wallace J, Barbian H, et al. CD32 is enriched on CD4dimCD8bright T cells. PLoS One. 2020;15(9):e0239157.

56. Zloza A, Sullivan YB, Connick E, Landay AL, Al-Harthi L. CD8+ T cells that express CD4 on their surface (CD4dimCD8bright T cells) recognize an antigen-specific target, are detected in vivo, and can be productively infected by T-tropic HIV. Blood. 2003;102(6):2156–2164.

57. Egawa T. A Fateful Decision in the Thymus Controlled by the Transcription Factor ThPOK. J.I. 2021;206(9):1981–1982.

58. Overgaard NH, Jung J-W, Steptoe RJ, Wells JW. CD4+/CD8+ double-positive T cells: more than just a developmental stage? J. Leukoc. Biol. 2015;97(1):31–38.

59. Hess NJ, Hudson AW, Hematti P, Gumperz JE. Early T Cell Activation Metrics Predict Graft-versus-Host Disease in a Humanized Mouse Model of Hematopoietic Stem Cell Transplantation. J.Immunol. 2020;205(1):272–281.

60. Hess NJ, S Bharadwaj N, Bobeck EA, et al. iNKT cells coordinate immune pathways to enable engraftment in nonconditioned hosts. Life Sci Alliance. 2021;4(7):.

61. Walsh AJ, Mueller K, Tweed K, et al. Classification of T-cell activation via autofluorescence lifetime imaging. Nat Biomed Eng. 2021;5(1):77–88.

62. Samimi K, Contreras Guzman E, Trier SM, et al. Time-domain single photon-excited autofluorescence lifetime for label-free detection of T cell activation. Opt Lett. 2021;46(9):2168–2171.

63. Hess NJ, Lindner PN, Vazquez J, et al. Different Human Immune Lineage Compositions Are Generated in Non-Conditioned NBSGW Mice Depending on HSPC Source. Front Immunol. 2020;ll:573406.

64. Reis BS, Rogoz A, Costa-Pinto FA, Taniuchi I, Mucida D. Mutual expression of the transcription factors Runx3 andThPOK regulates intestinal CD4^+^T cell immunity. Nat Immunol. 2013;14(3):271–280.

65. Mestas J, Hughes CCW. Of mice and not men: differences between mouse and human immunology. J. Immunol. 2004;172(5):2731–2738.

