## Supplemental Data for "Inflammatory CD4/CD8 double positive human T cells arise from reactive CD8 T cells and are sufficient to mediate GVHD pathology"

**Supplemental Methods**

*Two-photon lifetime imaging of NAD(P)H and FAD*

A custom-built inverted multiphoton microscope (Bruker Fluorescence Microscopy, Middleton, WI, USA), was used to acquire fluorescence intensity and lifetime images. The equipment consists of an ultrafast laser (Spectra Physics, Insight DSDual), an inverted microscope (Nikon, Eclipse Ti), and a 40X water immersion (1.15 NA, Nikon) objective. NAD(P)H and FAD images were obtained for the same field of view. FAD fluorescence was isolated using an emission bandpass filter of 550/100 nm and excitation wavelength of 890 nm. NAD(P)H fluorescence was isolated using an emission bandpass filter of 440/80 nm and an excitation wavelength of 750 nm. Fluorescence lifetime images were collected using time-correlated single-photon counting electronics (SPC-150, Becker and Hickl, Brookline, MA, USA) and a GaAsP photomultiplier tube (H7422P-40, Hamamatsu, Bridgewater, NJ, USA). 512x512 pixel images were obtained using a pixel dwell time of 4.8 µs over 60 s total integration time. To guarantee adequate photon observations for lifetime decay fits and no photobleaching, the photon count rates were maintained at 1–2 x 10^5^ photons/s. The instrument response function was calculated from the second harmonic generation of urea crystals excited at 900 nm. A Fluoresbrite YG microsphere (Polysciences Inc., Warrington, PA, USA) was imaged as a daily standard for fluorescence lifetime. The lifetime decay curves for the YG microsphere standard were fit to a single exponential decay and the fluorescence lifetime was measured to be 2.1 ns (n = 7), which is consistent with published values. NAD(P)H and FAD intensity and lifetime images were analyzed using SPCImage software (Becker &Hickl, Berlin, Germany) as described previously^65^. The fluorescence lifetime decay curve was deconvolved with the instrument response function and fit to a two-component exponential decay model at each pixel, I(t) = α_1*_ e^(-t/τ_1_) + α_2_*e^(-t/τ_2_) + C, where I(t) represents the fluorescence intensity at time t after the laser excitation pulse, α accounts for the fractional contribution from each component, C represents the background light, and τ is the fluorescence lifetime of each component. Since both NAD(P)H and FAD can exist in two conformational states, bound, or unbound to enzymes, a two-component model was used. For NAD(P)H, the short (τ_1_) and long (τ_2_) lifetime components correspond with the unbound and bound conformations, respectively. The mean lifetime (τ_m_) was calculated using τ_m_ = α_1*_ τ_1_+ α_2*_ τ_2_, for both NAD(P)H and FAD. The optical redox ratio was determined from the NAD(P)H and FAD intensity images as the intensity of NAD(P)H / (NAD(P)H + FAD). Semi-automated cell segmentation was performed using the cyto2 model in Cellpose, which resulted in whole-cell masks, followed by manual inspection and mask corrections by a trained scientist. Using the masks, single cell features were extracted including bi-exponential fit parameters (τ_1_, τ_2_, α_1_, α_2_) for NAD(P)H and FAD, NAD(P)H τ_m_, FAD τ_m_, and the optical redox ratio. Features were calculated on a per-cell level by averaging all the pixels across each cell. A minimum of 100 cells were analyzed per condition to obtain statistically robust results.

*UMAP Clustering*

Clustering of cells across all cell types (CD4, CD8 and DPT) of combined control and activated conditions was represented using UMAP. UMAP dimensionality reduction was performed on all 10 OMI variables (NAD(P)H τm, τ1, τ2, α1, α2, and FAD τm, τ1, τ2, α1, α2) for projection in 2D space. The following parameters were used for UMAP visualizations: “n _neighbors”: 10; “min_dist”: 0.5, “metric”: euclidean, “n_components”: 2.

*Classification Methods*

Random forest classifiers were trained to distinguish DPT cells from the combined group of CD4 and CD8 T cells. The DPT OMI data was randomly partitioned into training and validation datasets at the proportions of 70% and 30%, respectively (n = 179 cells in the training set, n = 419 cells in the validation set). Variable weights for OMI variables were extracted to determine the contribution of each variable to the trained random forest model: NAD(P)H τm = 0.07, τ1 = 0.19, τ2 = 0.09, α1 = 0.08, α2 = 0.08, and FAD τm = 0.12, τ1 = 0.09, τ2 = 0.11, α1 = 0.08, α2 = 0.09. Receiver operating characteristic (ROC) curves were generated to evaluate the random forest model performance on classification of test set data.

**Supplemental Tables**

**Supplemental Table 1. Patient Transplant Characteristics**

Summary of the transplant metrics from the 35 patients included in the observational study with the number and percentage for each variable. Patients were included in the study irrespective of their conditioning regimen, donor matching status, graft source, GVHD prophylaxis regimen or malignant disease type. All patients received post-transplant cyclophosphamide at day +3/4 (standard at our center). Patients were grouped into categories based on the maximum GVHD score in their first 100 days after transplant or a relapse event within 1 year of transplant.

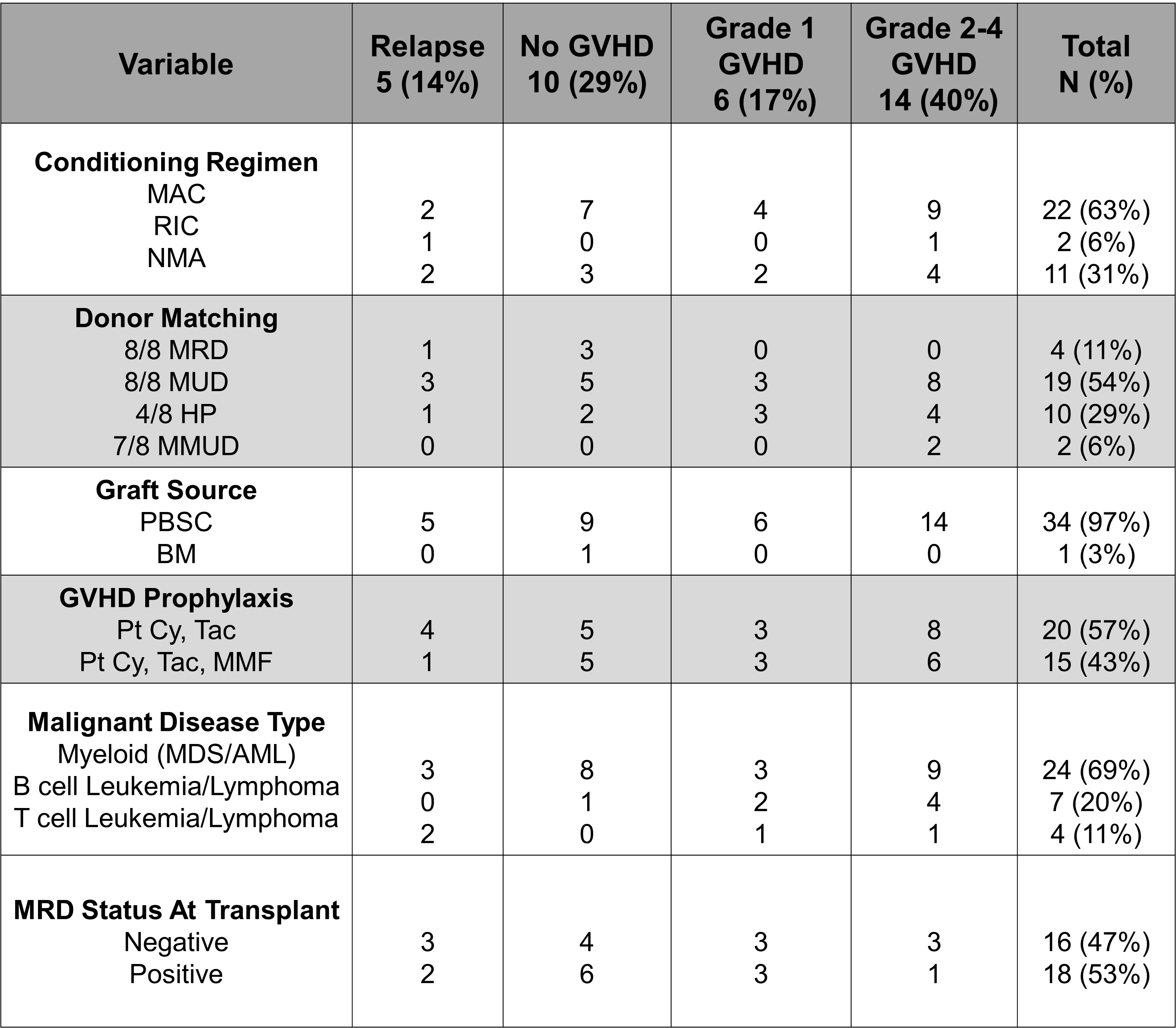

**Supplemental Table 2. Functional Hierarchy of Differentially Expressed Genes in DPTs.**

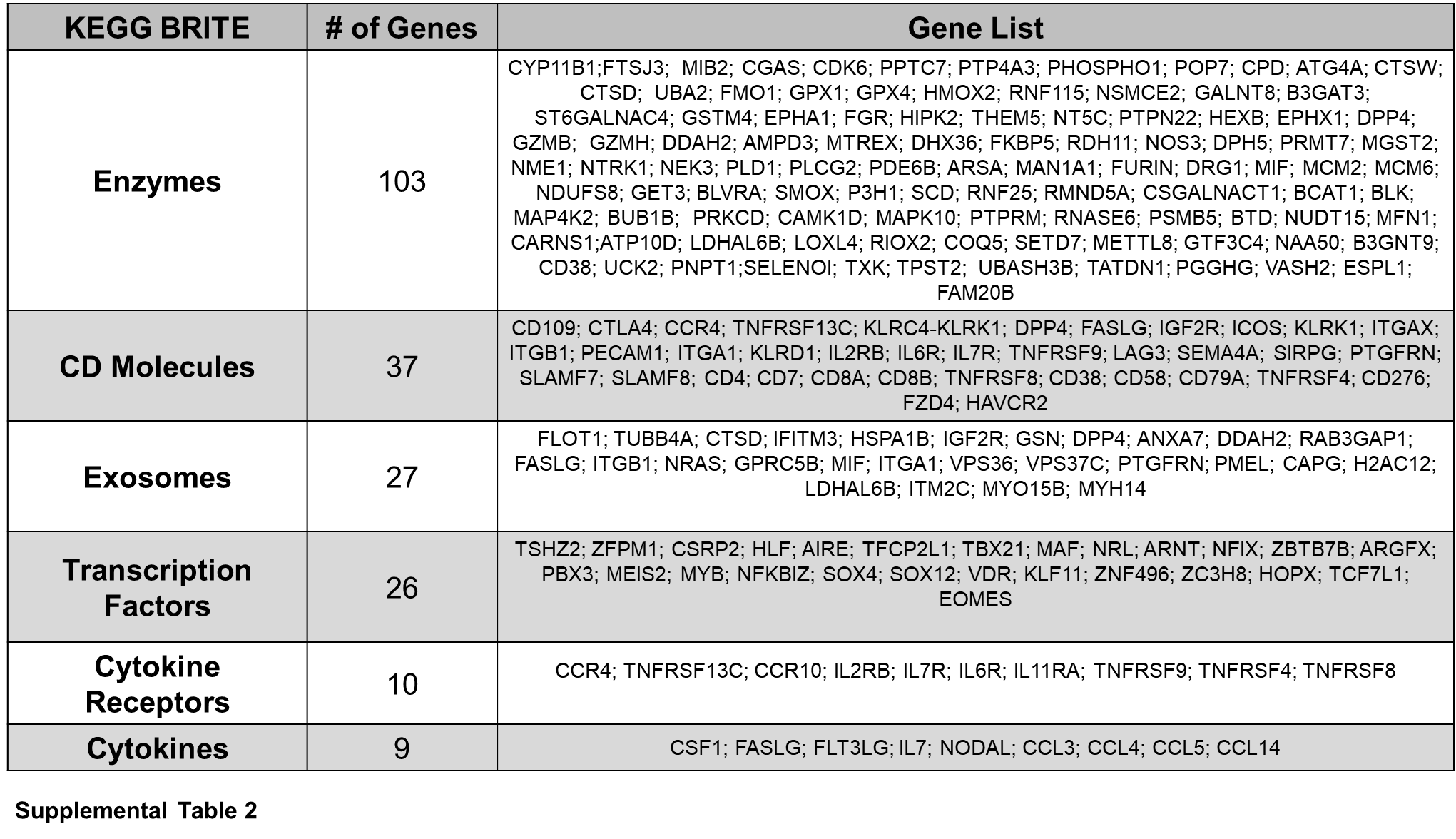
A combined total of 796 differentially expressed genes from the DPT:CD4 and DPT:CD8 analyses were analyzed using the KEGG BRITE database for functional hierarchical classification. Six BRITE classifications, the number of genes contributing to the classification and each respective gene name within the classification are listed.

**Supplemental Table 3. Clonotypes of Human T cells In Xenogeneic GVHD.**

List of the TRBV, -D, and -J gene segments associated with the ten highest clones identified by RNA-seq in the CD4, CD8 and DPT populations from xenogeneic GVHD mice.

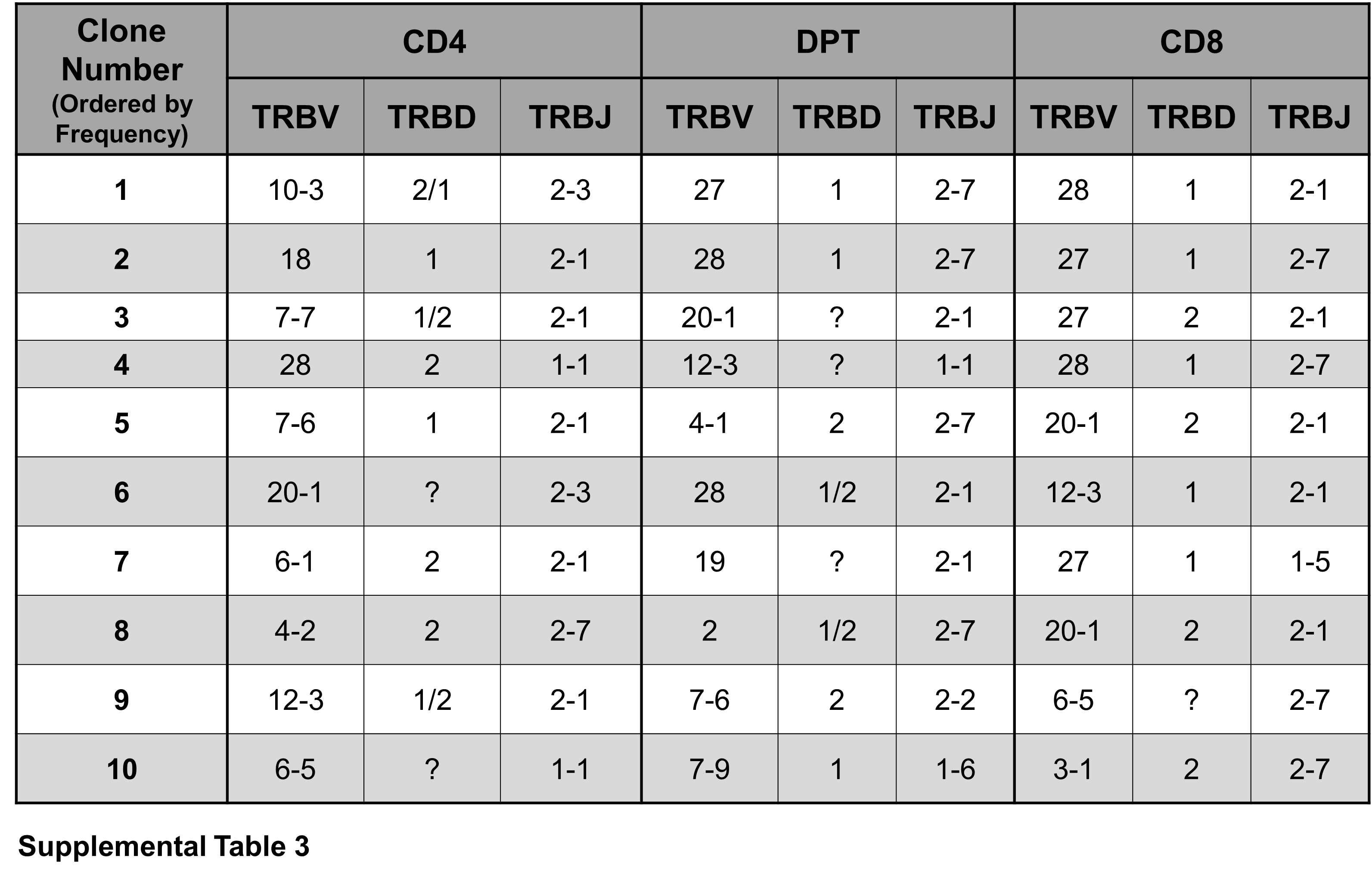

**Supplemental Figures**

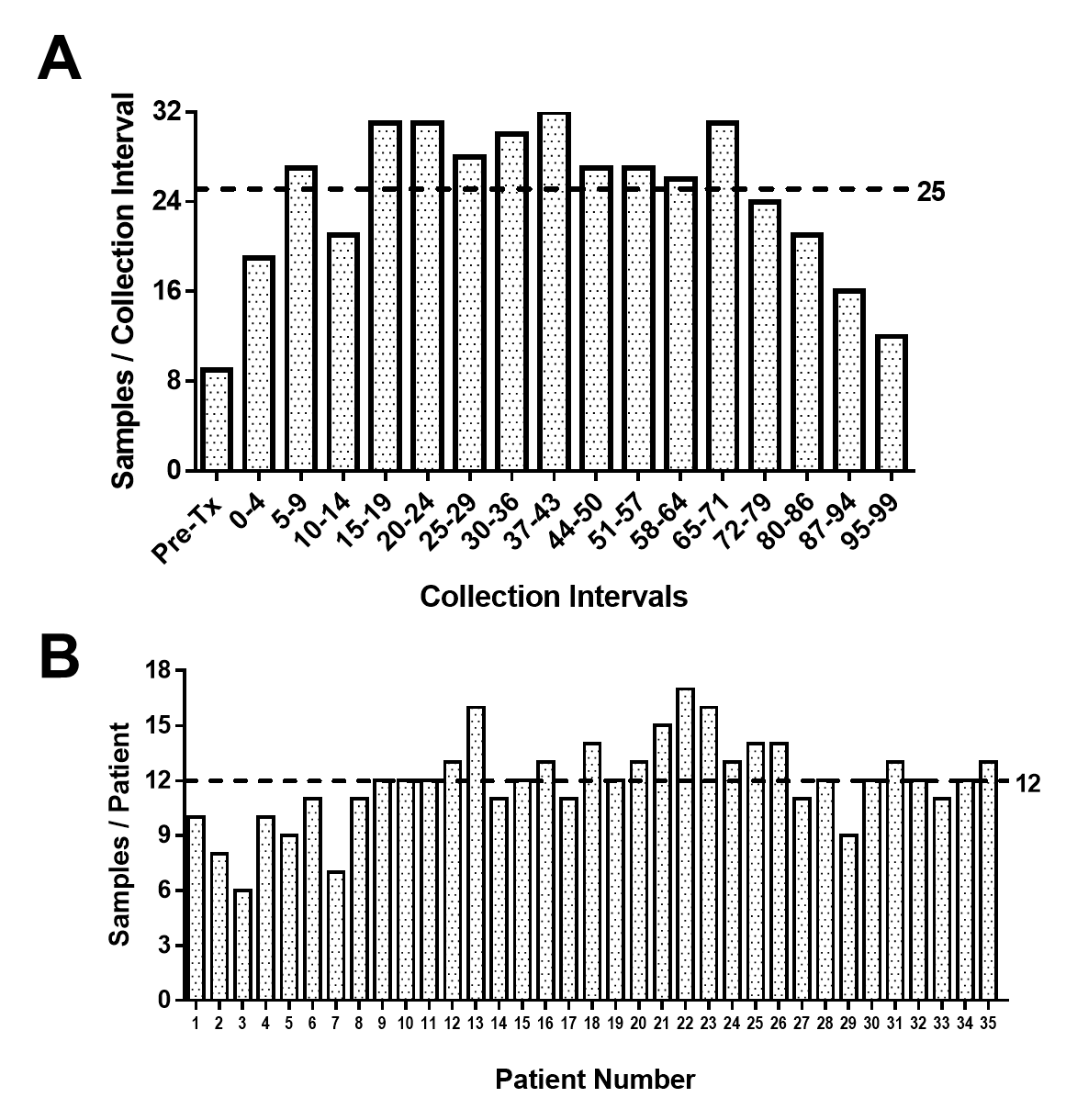

**Supplemental Figure 1. Sample collection across patients and time.**

The number of samples collected within each collection interval (A) and from each patient (B) are shown with the dashed line representing the average from our cohort.

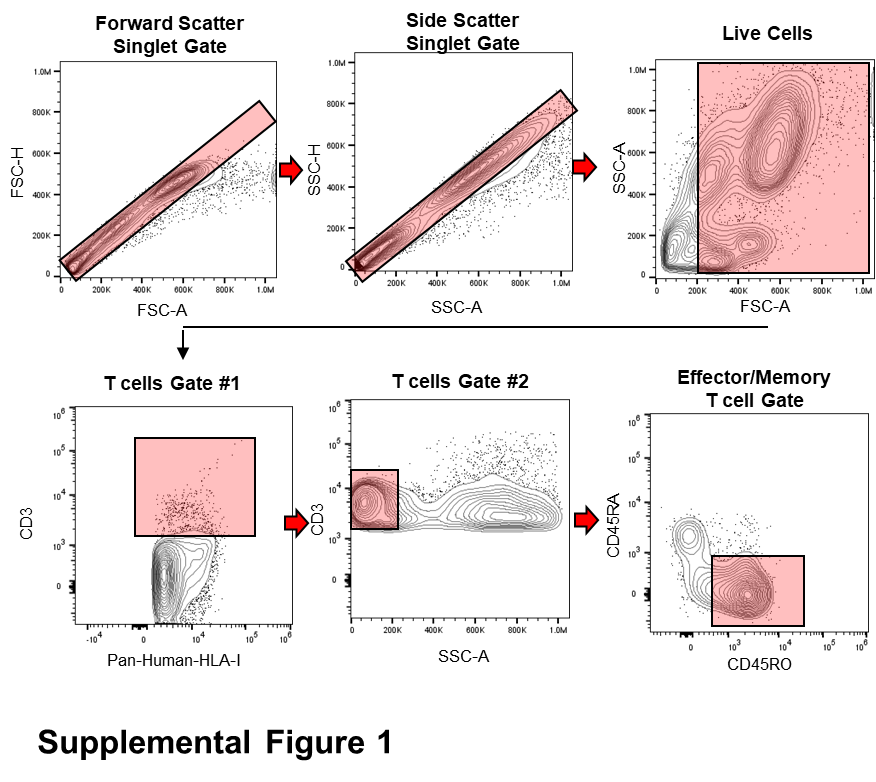

**Supplemental Figure 2. Flow cytometric gating strategy of clinical samples.**

Sequential gating strategy that was used on all patient samples across all the collection intervals. Red boxes denote gated populations. Forward and side scatter singlets were first gated followed by a live cell gate based on the leukocyte population from healthy peripheral blood samples. T cells were gated by pan-HLA-I and CD3 positive staining and T cells with excessive SSC-A were further removed as potential dead/dying cells. Effector/memory T cells were then gated as CD45RO positive cells.

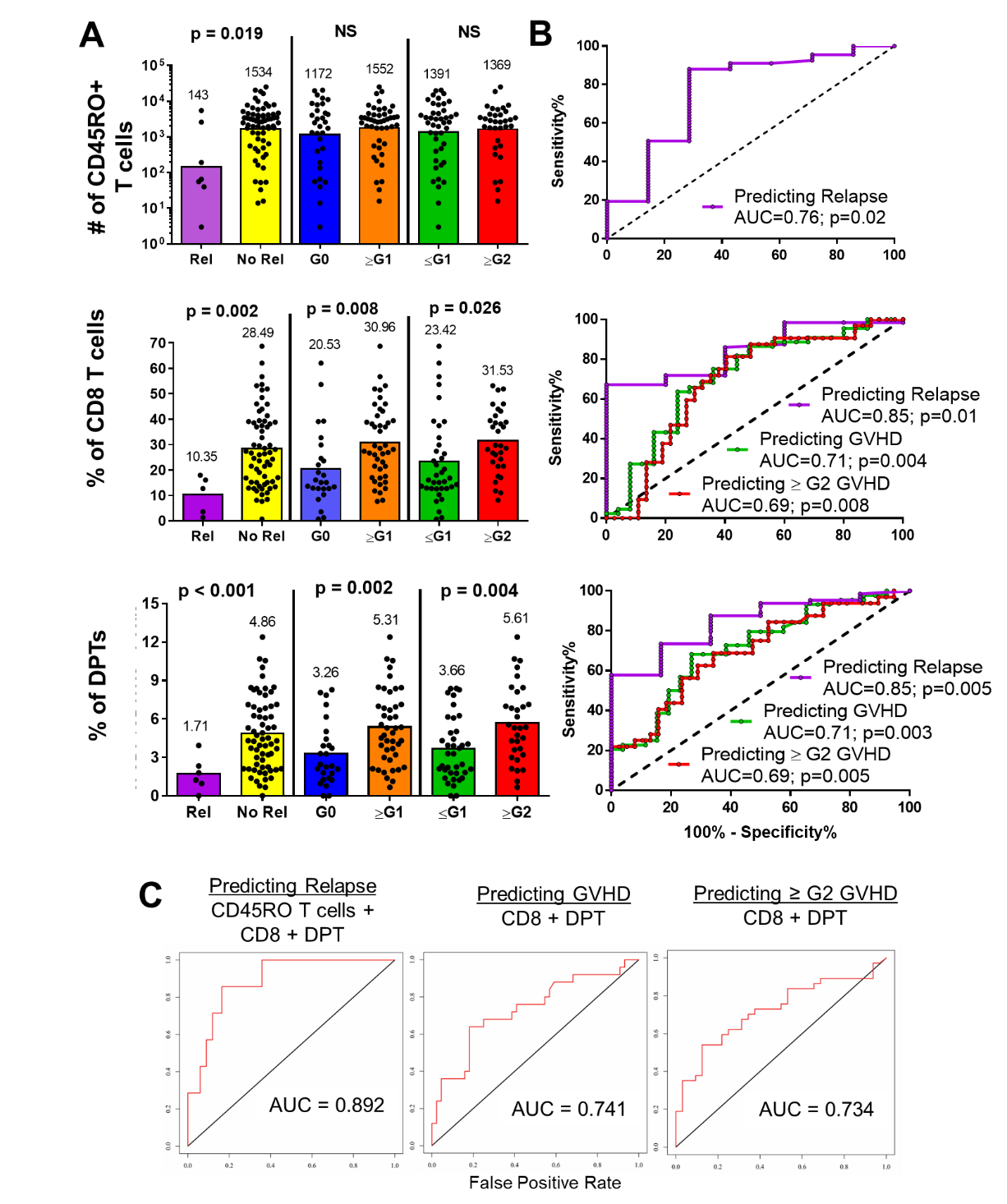

**Supplemental Figure 3. T cell metrics collected after day 22 can predict allo-HSCT outcomes**

Blood samples from allo-HSCT recipients were collected and analyzed as described in Fig 1. Data represents samples collected between 22- and 83-days post-transplant. Samples were further normalized to the date of each patient’s G1 or G2 GVHD diagnosis (whichever was higher). Patients who did not develop GVHD were normalized to day 54 which was the median time to GVHD diagnosis in our cohort. Samples from the patient’s date of diagnosis and two collection intervals prior were included in this analysis. As in Fig 1, bar graphs of the three binary tests are shown (A) with tests reaching a significance of 0.05 further analyzed by ROC (B). T cell metrics predicting the same allo-HSCT outcome were combined in a multi-parameter ROC analysis shown in (C).

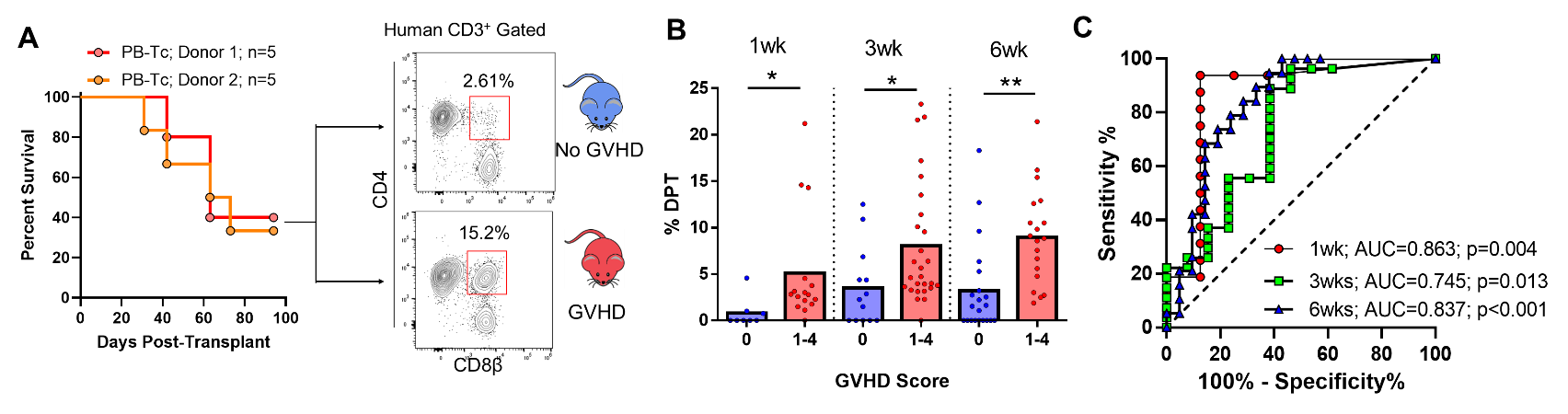

**Supplemental Figure 4. Isolated human T cell grafts develop DPTs after transplantation**

(A) Survival curve of NSG mice transplanted with isolated human PB-T cells from two different donors. Mice were divided into a GVHD and No GVHD group based on lethality by 12 weeks post-transplant. Representative dot plot of the human T cell population taken from the blood at 3wks post-transplant for each group is also shown. (B) Graphical representation of the percentage of DPTs in the blood of mice at 1-, 3- and 6-weeks post-transplant. (C) ROC analysis from the data shown in (B) with the AUC and significance values indicated. PB = peripheral blood. * p < 0.05, ** p < 0.01.

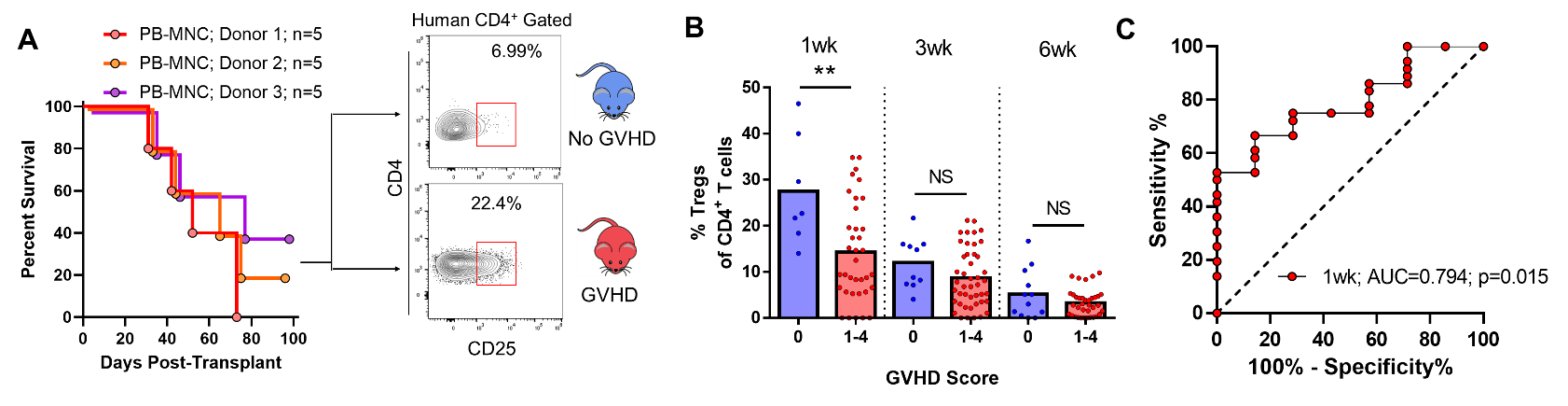

**Supplemental Figure 5. Tregs are predictive of lethal GVHD after xenogeneic transplantation.**

(A) Survival curve of NSG mice transplanted with isolated human PB-MNC from three different donors. Mice were divided into a GVHD and No GVHD group based on lethality by 12 weeks post-transplant. Representative dot plot of the human T cell population taken from the blood at 3wks post-transplant for each group is also shown. (B) Graphical representation of the percentage of Tregs in the blood of mice at 1-, 3- and 6-weeks post-transplant. (C) ROC analysis from the data shown in (B) with the AUC and significance values indicated. PB = peripheral blood, NS = not significant. * p < 0.05, ** p < 0.01.

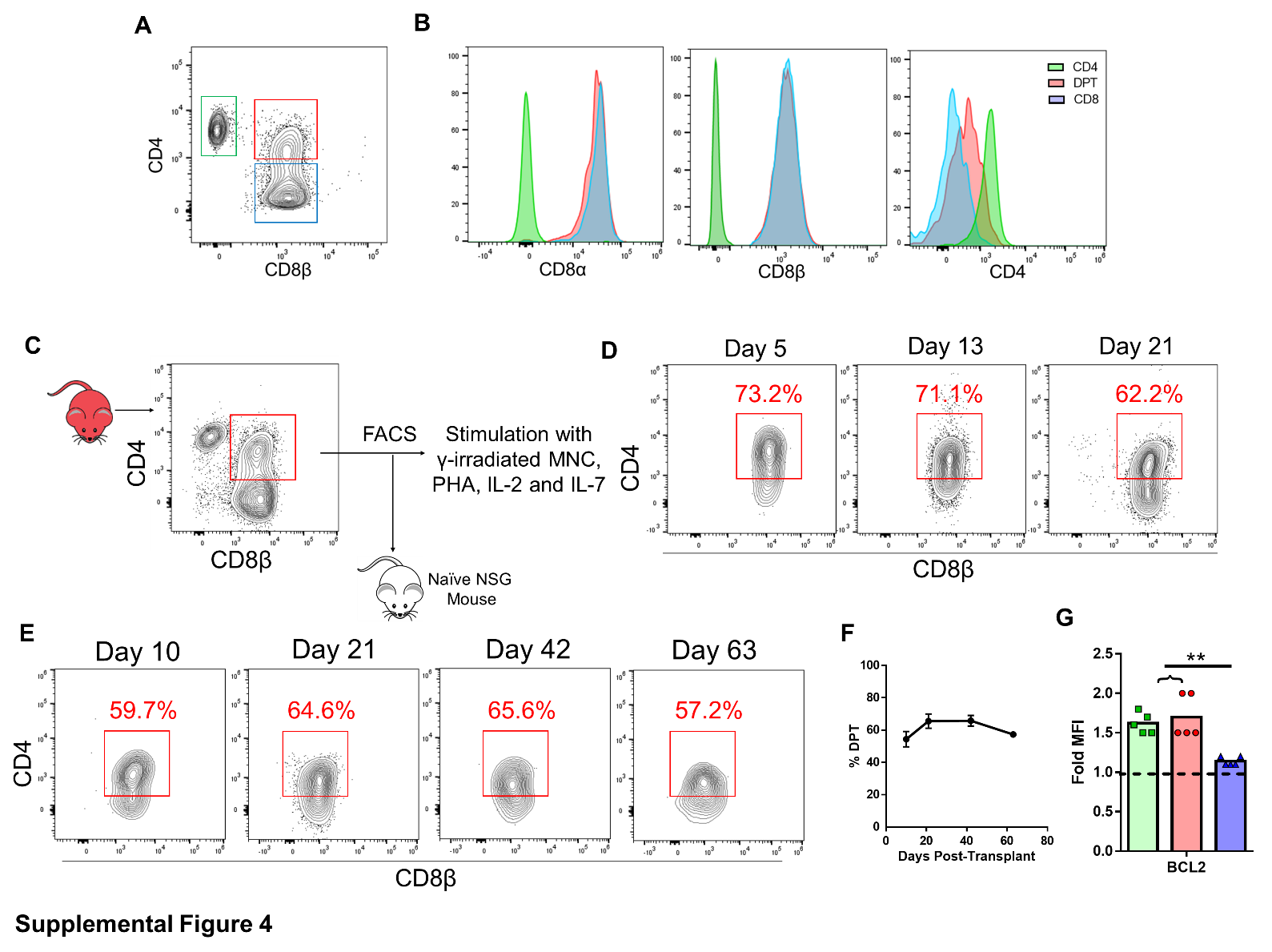

**Supplemental Figure 6. Expression of CD8 Is Maintained In DPTs Following in vitro or in vivo Stimulation**

(A) Gating strategy used for CD4 (green), DPTs (red) and CD8 T cells (blue) isolated from a xeno-GVHD mouse. (B) Representative histograms of CD8α, CD8β and CD4 expression on each T cell population gated in (A). (C) DPTs (red box) were flow sorted and then either expanded in vitro using γ-irradiated MNC, IL-2 and IL-7 for three weeks (D) or re-transplanted back into a naïve mouse (E-F). Representative dot plots of DPTs are shown (D-E) with the percentage of DPTs in the blood of mice quantified from five mice in (F). (G) Fold MFI of the pro-survival protein BCL2 as measured by intracellular flow cytometry. Each dot represents an individual mouse. Parametric t-test was used to determine significance. ** p<0.01.

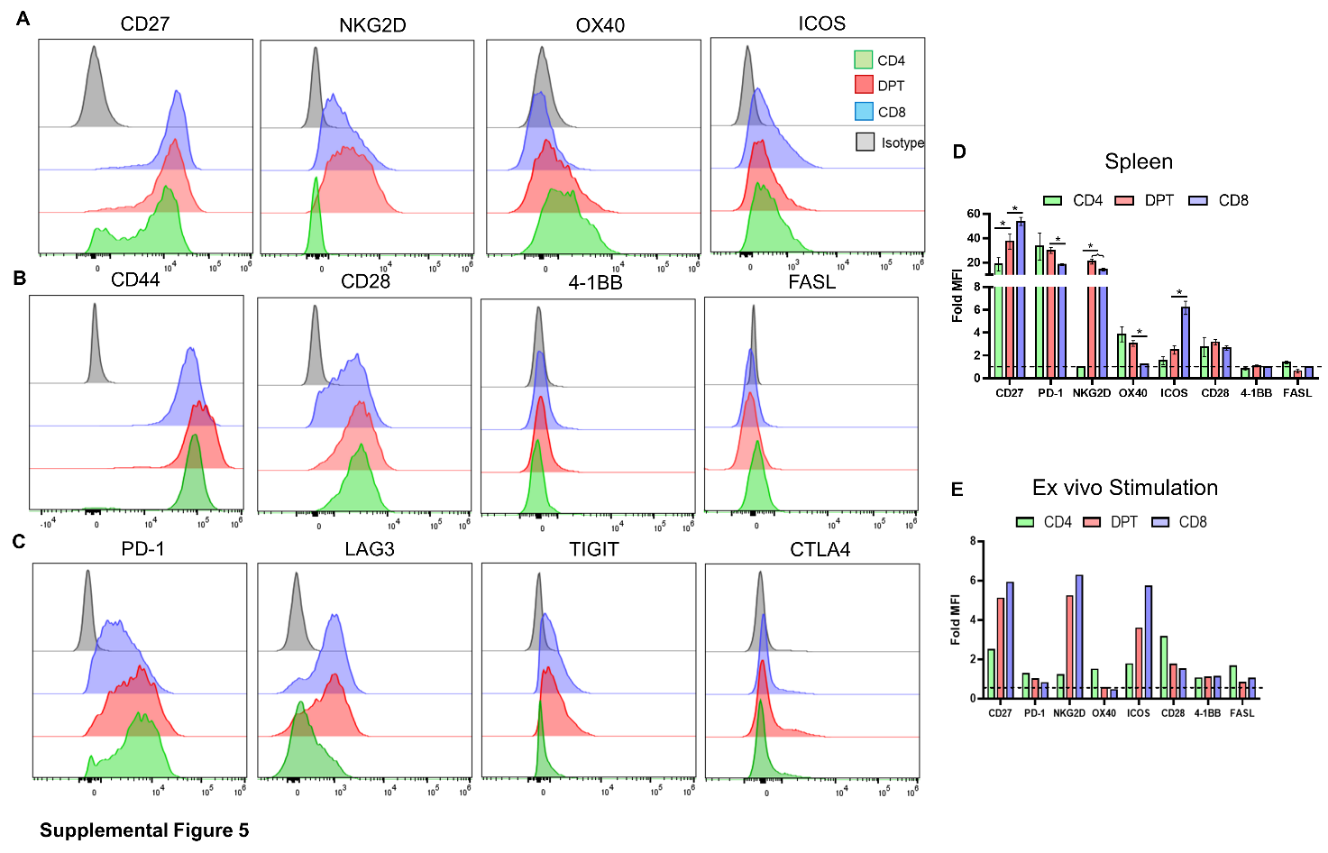

**Supplemental Figure 7. DPT display expression markers of both CD4 and CD8 T cells.**

(A) Representative histograms of the expression of 12 different markers on the surface of CD4, DPT and CD8 T cells from the blood of xeno-GVHD mice. Histograms are divided by pro-inflammatory markers having differential expression between T cell populations (A), pro-inflammatory markers without any expression differences (B) and inhibitory markers (C). (D-E) Graphical representation of eight different markers analyzed from T cells taken from the spleens of xeno-GVHD mice (D) or ex vivo stimulated as in Supplemental Figure 2C-D (E). Parametric t-test was used to determine significance. * p<0.05

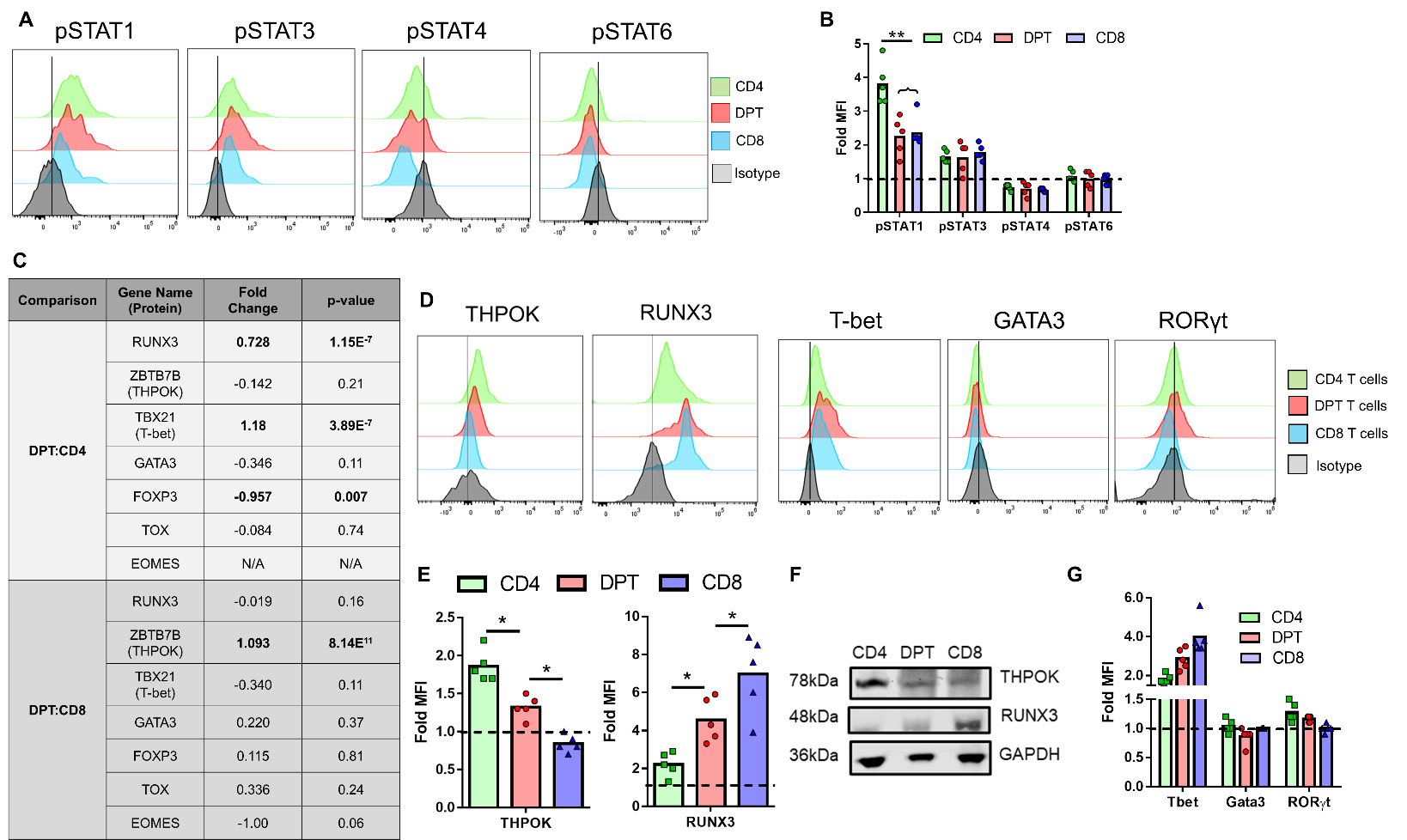

**Supplemental Figure 8. DPTs do not display a distinct STAT or transcription factor profile.**

(A-B) Representative histograms and graphical analysis of the Fold MFI of the indicated phospho-STAT proteins of T cells from the blood of xeno-GVHD mice. (C) Table highlighting the fold change and p-value of a subset of relevant T cell transcription factors from our RNA-seq dataset comparing DPT to CD4 and CD8 T cells. (D-G) Representative histograms (D) and graphical analysis (E, G) and western blot data (F) of five T cell transcription factors. All data is representative of at least two independent experiments containing at least five mice each. Each dot represents an individual mouse. Parametric t-test was used to determine significance. * p<0.05, ** p<0.01

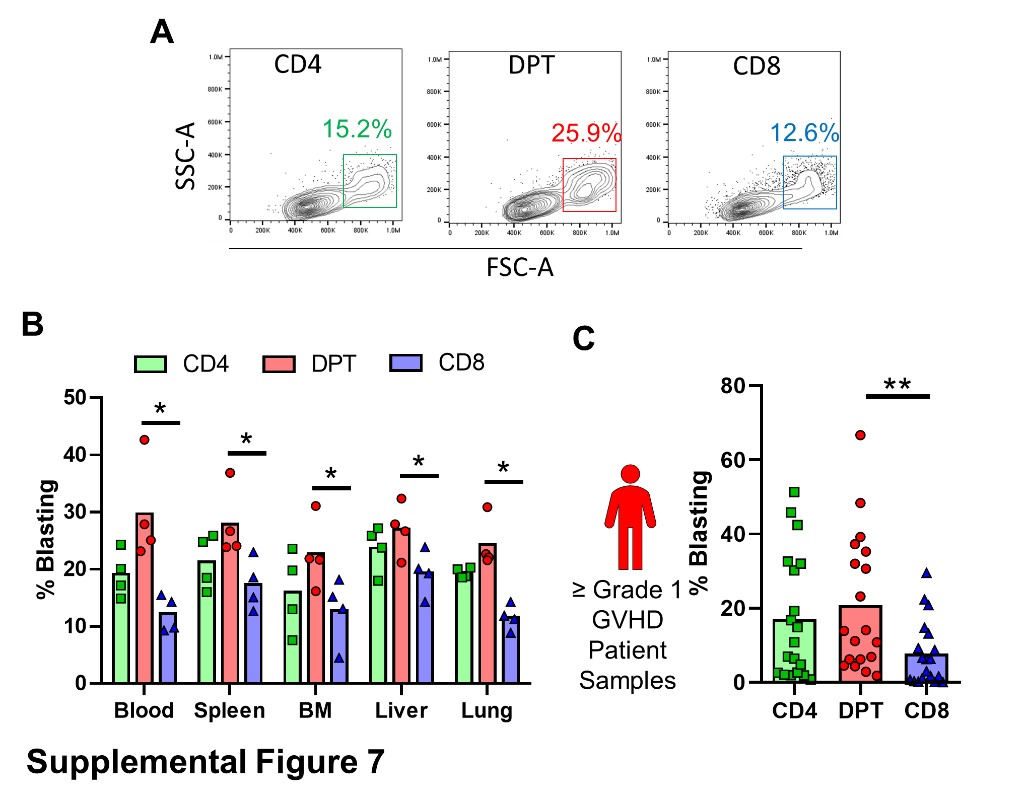

**Supplemental Figure 9. DPTs display a phenotype of proliferating T cells.**

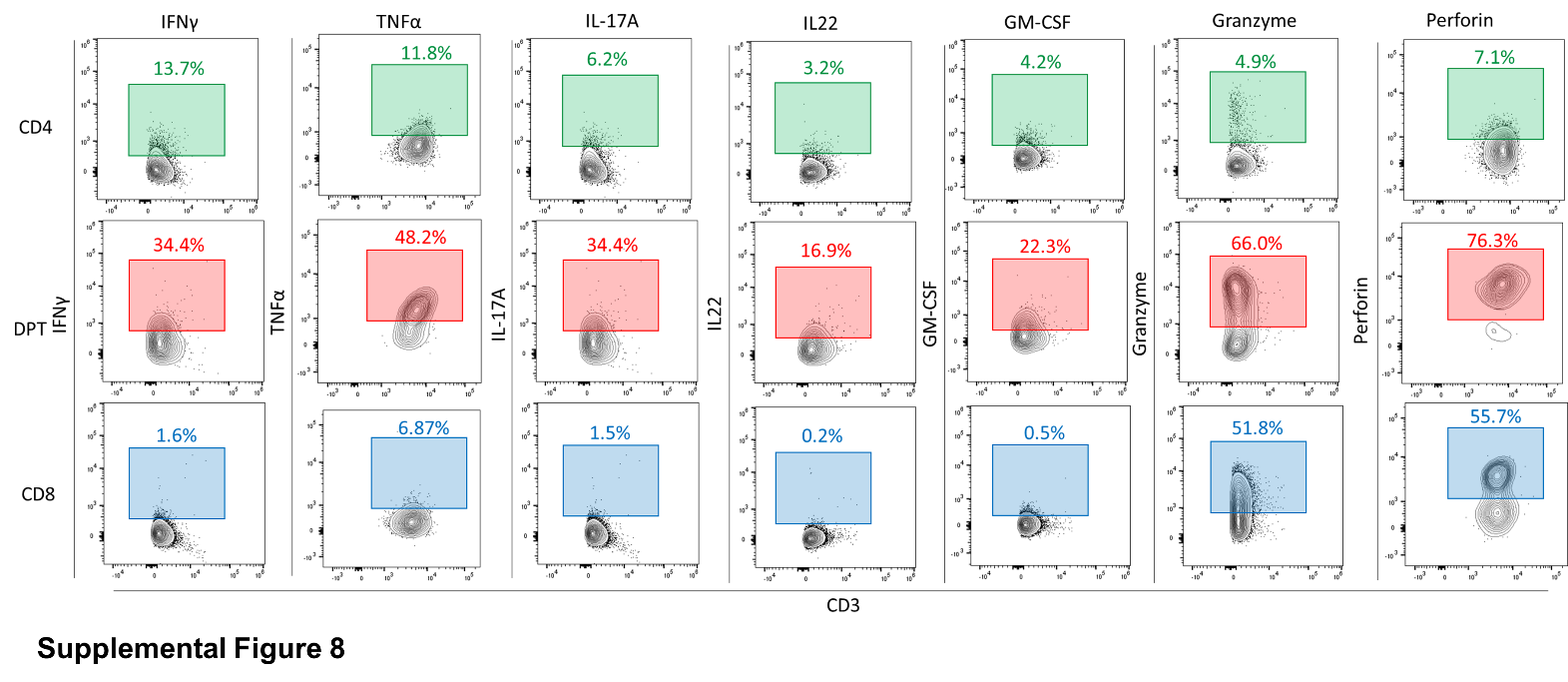
(A) Representative dot pot, gating strategy and percentage of blasting T cells from the blood of xeno-GVHD mice taken at 6-weeks post-transplant. (B) Graphical representation of (A) including data from additional xeno-GVHD target organs. (C) Percentage of blasting cells within the CD4, DPT and CD8 populations taken from 19 of the allo-HSCT patients in our observational study at the time of their GVHD diagnosis. Parametric t-test was used to determine significance. * p<0.05, ** p<0.01, *** p<0.001

**Supplemental Figure 10. DPTs express both cytolytic and modulatory cytokines**

T cells from the blood of xeno-GVHD mice were incubated overnight with either brefeldin A alone or brefeldin A followed by a four-hour PMA/ionomycin stimulation. Seven inflammatory cytokines were measured by ICS from between 4-13 xeno-GVHD mice with the gating strategy and percentage shown for each cytokine.

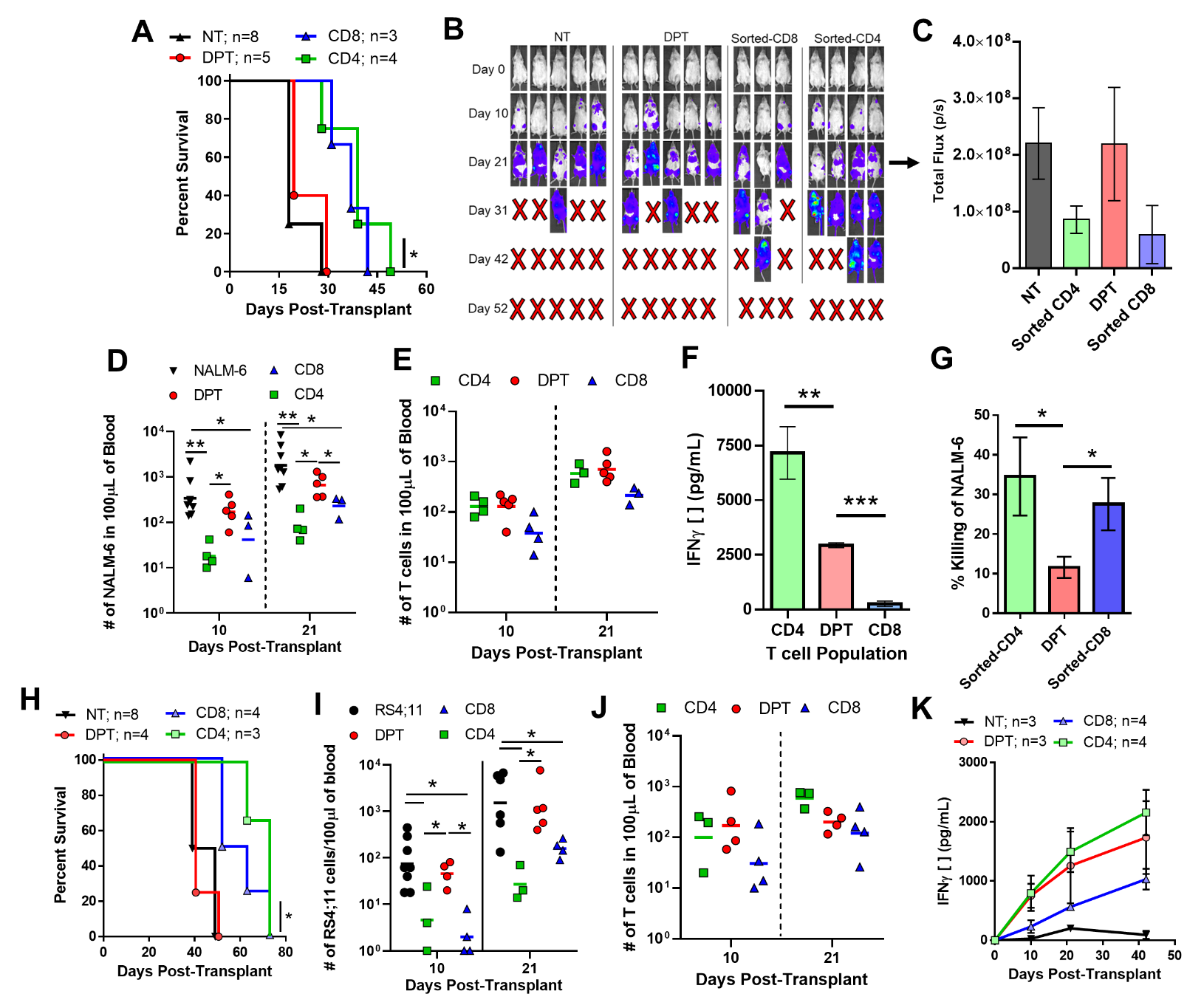

**Supplemental Figure 11. Freshly isolated CD4 and CD8 T cells display similar GVL activity compared to CD4 and CD8 T cells sorted from GVHD mice.**

As in Fig 8, mice were first transplanted with the B-ALL cell line NALM-6 (A-G) or RS4;11 (H-K) prior to transplantation with human CD4, DPTs or CD8 T cells sorted from xeno-GVHD mice. Survival curve (A), IVIS images (B), IVIS quantification (C), number of NALM-6 (D) and T cells (E) from the blood xeno-GVHD mice at 10- and 21-days post-transplant. (E) IFNγ from the plasma was also collected at 3 weeks post-transplant. (G) As in Fig 8H, NALM-6 were pre-loaded with Calcein prior to co-culture with CD4, DPT or CD8 T cells sorted from xeno-GVHD mice for four hours and the supernatant analyzed for Calcein fluorescence. Data is shown relative to max killing with Triton-X. Data from (A-G) includes DPT data also shown in Fig 8B-H for comparison purposes. (H-K) Experiment was repeated using the human B-ALL cell line RS4;11 with the survival curve (H), number of RS4;11 (I) and T cells (J) shown along with IFNγ quantification (K). Non-parametric t-tests were used to determine significance of total NALM-6 and T cells in the blood of mice, parametric t-tests were used for all other tests. * p<0.05, ** p<0.01
